# Promoter scanning during transcription initiation in *Saccharomyces cerevisiae*: Pol II in the “shooting gallery”

**DOI:** 10.1101/810127

**Authors:** Chenxi Qiu, Huiyan Jin, Irina Vvedenskaya, Jordi Abante Llenas, Tingting Zhao, Indranil Malik, Alex M. Visbisky, Scott L. Schwartz, Ping Cui, Pavel Čabart, Kang Hoo Han, William K. M. Lai, Richard P. Metz, Charles D. Johnson, Sing-Hoi Sze, B. Franklin Pugh, Bryce E. Nickels, Craig D. Kaplan

## Abstract

**Background:** The majority of eukaryotic promoters utilize multiple transcription start sites (TSSs). How multiple TSSs are specified at individual promoters across eukaryotes is not understood for most species. In *S. cerevisiae*, a preinitiation complex comprised of Pol II and conserved general transcription factors (GTFs) assembles and opens DNA upstream of TSSs. Evidence from model promoters indicates that the preinitiation complex (PIC) scans from upstream to downstream to identify TSSs. Prior results suggest that TSS distributions at promoters where scanning occurs shift in a polar fashion upon alteration in Pol II catalytic activity or GTF function.

**Results:** To determine extent of promoter scanning across promoter classes in *S. cerevisiae*, we perturbed Pol II catalytic activity and GTF function and analyzed their effects on TSS usage genome-wide. We find that alterations to Pol II, TFIIB, or TFIIF function widely alter the initiation landscape consistent with promoter scanning operating at all yeast promoters, regardless of promoter class. Promoter architecture, however, can determine extent of promoter sensitivity to altered Pol II activity in ways that are predicted by a scanning model.

**Conclusions:** Our observations coupled with previous data validate key predictions of the scanning model for Pol II initiation in yeast – which we term the “shooting gallery”. In this model, Pol II catalytic activity, and the rate and processivity of Pol II scanning together with promoter sequence determine the distribution of TSSs and their usage.

## BACKGROUND

Gene expression can be regulated at all levels, and its proper control is critical for cellular function. Transcription regulation has been of intense interest for decades as it determines how much RNA is synthesized for a given gene or locus. Much regulation occurs at the first step in transcription, initiation. A multitude of signals can be integrated with the activities of transcriptional regulators that converge on individual gene promoters. Subsequent to the integration of regulatory information, RNA Polymerase II (Pol II) and general transcription factors (GTFs) must recognize core promoters to together initiate transcription at specific sequences, transcription start sites (TSSs). As with any biochemical process, the efficiency of individual steps will shape the overall output. Thus, determinants of core promoter output during initiation, both overall expression level and the exact position of transcription start sites (TSSs), will be affected by the efficiency of biochemical events during initiation. How different core promoters modulate biochemical steps in initiation, and the nature of their functional interactions with the initiation machinery, are not fully understood.

Classes of eukaryotic core promoters can be distinguished by DNA sequence motifs and chromatin structure (reviews of the core promoter over time [1-10]). These features together comprise a promoter’s architecture, which may also correlate with differential recruitment or requirement for particular GTF complexes [11-13]. A theme across eukaryotes is that core promoters can be broadly separated into two main classes by examination of architectural features and factor requirements. A number of studies indicate that the most common eukaryotic promoters are nucleosome-depleted regions (NDRs) flanked by positioned nucleosomes, which can support divergent transcription through assembly of pre-initiation complexes (PICs) proximal to flanking nucleosomes (with exceptions)[14-25]. We will adhere to the definition of “core promoter” as representing the DNA elements and chromatin structure that facilitate transcription in one direction, to avoid definitional confusion that a “promoter” inherently drives divergent transcription [26-28]. In yeast, promoter classes have been distinguished in many ways with the end result generally being two main classes of promoter are recognized [16-18, 29, 30]. These classes are distinguished by the presence or absence of a consensus TATA element [31, 32], presence or absence of stereotypical nucleosome organization [18], enrichment for specific transcription factor binding [14, 33, 34], enrichment for non-TATA sequence motifs [35, 36], and differential sensitivity to mutations in GTFs or transcription coactivators [31, 33, 34]. Core promoters attached to defined NDRs tend to lack canonical TATA-elements. Conversely, in yeast and other eukaryotes, core promoters with TATA elements can lack stereotypical nucleosome organization and may have nucleosomes positioned over the TATA box in the absence of gene activation. While there have been a number of additional core promoter elements identified in other organisms, especially *Drosophila melanogaster* [37], we will focus on the distinction provided by presence or absence of TATA-elements.

The TATA element serves as a platform for core promoter binding of the TATA-Binding Protein (TBP). TBP recognition of promoter DNA is assumed to be critical for PIC formation and Pol II promoter specificity. Functional distinction in promoter classes is supported by studies showing differential factor recruitment and requirements between them, with TATA promoters showing higher SAGA dependence and reduced Taf1 (a TFIID subunit) recruitment [31-34], though recent data have been interpreted as both SAGA and TFIID functioning at all yeast promoters [38, 39], a distinction between the two classes seems to hold [40]. Conversely, TATA-less promoters show higher Taf1 recruitment and greater requirement for TBP-Associated Factor (TAF) function. Given differences in reported factor requirements and promoter architectures, it is important to understand the mechanistic differences between promoters and how these relate to gene regulation.

TSS selection in *Saccharomyces cerevisiae* has been used as a model to understand how initiation factors collaborate to promote initiation [41, 42]. The vast majority of yeast core promoters specify multiple TSSs [43-45], and multiple TSS usage is now known to be common to the majority of core promoters in other eukaryotes [46-50]. Biochemical properties of RNA polymerase initiation lead to TSSs selectively occurring at a purine (R=A or G) just downstream from a pyrimidine (Y=C or T) – the Y_-1_R_+1_ motif [51]. Y_-1_R_+1_ motifs may be additionally embedded in longer sequence motifs (the Inr element)[52, 53]. In yeast, the initiation factor TFIIB has been proposed to “read” TSS sequences to promote recognition of appropriate TSSs, with structural evidence supporting positioning of TFIIB to read DNA sequences upstream of a TSS [11, 54].

Yeast differs from other model eukaryotes in that TSSs for TATA-containing core promoters are generally dispersed, and are found ∼40-120 nt downstream from the TATA [55]. Conversely, TSSs at TATA promoters in other organisms are tightly associated ∼31 nt downstream of the TATA (with the first T in “TATA” being +1)[56]. As TATA promoters represent ∼10% of promoters across well-studied organisms, they are the minority. Classic experiments using permanganate footprinting of melted DNA showed that promoter melting at two TATA promoters in yeast, *GAL1* and *GAL10*, occurs far upstream of TSSs, at a distance downstream from TATA where melting would occur in other eukaryotes that have TSSs closer to the TATA element [57]. This discovery led Giardina and Lis to propose that yeast Pol II scans downstream from TATA boxes to find TSSs. A large number of mutants have been found in yeast that perturb TSS selection, allowing the genetic architecture of Pol II initiation to be dissected, from those in Pol II subunit encoding genes *RPB1, RPB2, RPB7*, and *RPB9*, to GTF encoding genes *SUA7* (TFIIB), *TFG1* and *TFG2* (TFIIF), and *SSL2* (TFIIH), and the conserved transcription cofactor *SUB1* [58-78]. Mutants in GTFs or Pol II subunits have been consistently found at model promoters to alter TSS usage distributions in a polar fashion by shifting TSS distributions upstream or downstream relative to WT. These observations coupled with analysis of TSS mutations strongly support the directional scanning model for Pol II initiation (elegantly formulated in the work of Kuehner and Brow)[61].

Previous models for how initiation might be affected by Pol II mutants suggested that Pol II surfaces important for initiation functioned through interactions with GTFs within the PIC. We have previously found that altering residues deep in the Pol II active site, unlikely to be directly interacting with GTFs, but instead altering Pol II catalytic activity, had strong, allele-specific effects on TSS selection for model promoters [79-81]. Observed effects were polar in nature, and consistent with the Pol II active site acting downstream of a scanning process but during TSS selection and not afterwards. In other words, Pol II catalytic efficiency appears to directly impact TSS selection. For example, it appeared that increased Pol II catalytic activity increased initiation probability, leading to an upstream shift in TSS usage at candidate promoters because less DNA needs to be scanned on average prior to initiation. Conversely, lowering Pol II catalytic activity results in downstream shifts to TSS usage at candidate promoters, because more promoter DNA has to be scanned prior to initiation. In general, candidate promoters examined for TSS selection have mostly been TATA containing (for example *ADH1, HIS4*), thus it is not known how universal Pol II initiation behavior or mechanisms are across all yeast core promoters, which likely comprise different classes with distinct architectures. To examine initiation by promoter scanning on a global scale in yeast, we perturbed Pol II or GTF activity genetically to examine changes to TSS usage across a comprehensive set of promoters that likely represent all yeast promoter classes. We have found that promoter scanning appears to be universal across yeast core promoters. Furthermore, we find that core promoter architecture correlates with sensitivity of core promoters to TSS perturbation in Pol II and initiation factor mutants. Our results have enabled formulation a model where Pol II and GTF function together in initiation to promote Pol II initiation efficiency at favorable DNA sequences. Finally, initiation by core promoter scanning prescribes a specific relationship between usable TSSs in a core promoter and the distribution of TSS usage, potentially allowing TSS distributions to be predicted if the sequence preferences for Pol II initiation can be measured.

## RESULTS

### Initiation mutants affect TSS selection globally in *Saccharomyces cerevisiae*

We previously found that yeast strains mutant for Pol II key active site residues important for normal catalysis showed polar effects on TSS selection at the model *ADH1* promoter in addition to some other promoters [80, 81]. *ADH1* is a TATA-containing promoter with major TSSs positioned at 90 and 100 nucleotides downstream of its TATA box. A number of mutants in Pol II and initiation factors also show TSS selection effects at *ADH1*. TSS selection effects have been hypothesized to relate to alterations in initiation sequence specificity, while the stereotypical polar effects of TSS-altering mutants are consistent with effects on scanning and not necessarily sequence specificity. These are not mutually exclusive models, and to understand better how Pol II activity and GTFs cooperate to identify TSSs, we mapped capped RNA 5′ ends genome-wide in *S. cerevisiae* using TSS-seq for WT, a series of Pol II catalytic mutants, a TFIIB mutant (*sua7-58A5*)[79], and a TFIIF mutant (*tfg2*Δ*146-180*)[82]. Positions of capped RNA 5′ ends are taken to represent positions of TSSs as Pol II-initiated RNA 5′ ends are capped shortly after emerging from the enzyme after initiation. We first determined how reproducible our pipeline (**Figure 1A**) was across the yeast genome, examining correlation of read positions corresponding to 5′ ends across all genome positions containing at least three mapped reads in each library being compared (**Figure 1B, Supplemental Figure 1A**). Examples of correlations between biological replicates are shown in **Figure 1B** for WT, one catalytically fast Pol II allele (*rpb1* E1103G)[83-85], and one catalytically slow Pol II allele (*rpb1* H1085Y)[81]. We refer to fast Pol II alleles as “gain of function” (GOF) alleles and slow Pol II alleles as “loss of function” (LOF) alleles [86]. Correlation plots for all other strains are shown in **Supplemental Figure 1A**. Clustering analysis of Pearson correlation coefficients among libraries aggregated from biological replicates for each strain indicates that Pol II and initiation mutant classes can be distinguished based on RNA 5′ end mapping alone (**Figure 1C**). **Supplemental Figure 1B** shows clustering of Pearson correlation coefficients of individual biological replicate TSS-seq libraries for reads within promoter regions.

**Figure 1.**
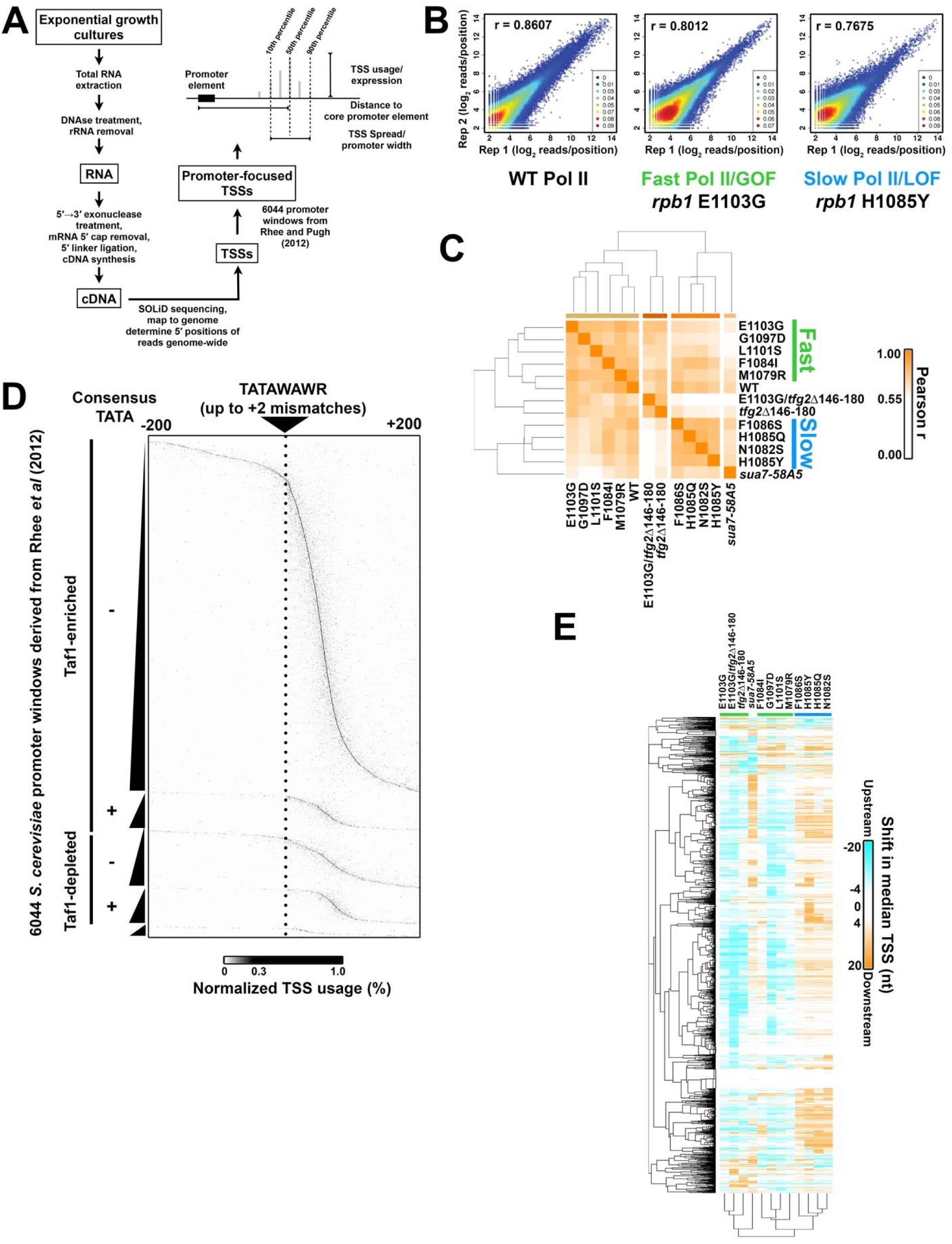
Genome-wide analysis of TSS selection in *S. cerevisiae*. **A.** Overview of method and description of simple metrics used in analyzing TSS distributions at yeast promoters. **B.** Reproducibility of TSS-seq analysis demonstrated by heat scatter correlation plots determine RNA 5′ ends across all genome positions with ≥3 reads in each library for biological replicates of WT, *rpb1* E1103G, and *rpb1* H1085Y libraries. Colors indicate the plot density from cool to warm (low to high estimated kernel density). Pearson r is shown. **C.** Heat map illustrating hierarchical clustering of Pearson correlation coefficients between aggregate (combined biological replicate) libraries for all strains. Clustering illustrates increased correlation among known reduced function *rpb1* alleles (“slow” or LOF) are increased correlation among increased activity *rpb1* alleles (“fast” or GOF). WT shows intermediate correlations with both classes. **D.** Core promoters (n=6044) predicted by Rhee and Pugh from GTF ChIP-exo data were used to initially map TSSs. TSSs were row normalized to illustrate the distribution within each window, TSSs generally map downstream of predicted core-promoters for most but not all promoter windows. Note that the resolution of the figure will have less pixels that promoter rows (6044). **E.** Determination of change in median TSS position (upstream shift in median position is negative (cyan), downstream shift in median position is positive (orange), see Methods) for promoters with ≥200 reads (n=3494). Heat map shows individual yeast promoter regions hierarchically clustered on the *y*-axis with the measured TSS shift for hierarchically-clustered TSS-usage affecting mutants on the *x*-axis. =

We first focused our analyses on promoter windows predicted from the localization of PIC components by Rhee and Pugh [14] and anchored on TATA or “TATA-like” elements (core promoter elements or CPE underlying PIC assembly points) at the +1 position of the promoter window (**Figure 1D**). RNA 5′ ends mapping to the top genome strand of these putative promoter windows indicates that these windows are associated with putative TSSs as expected. The majority of observed TSSs are downstream of predicted CPE/PIC locations from Rhee and Pugh, with TSSs originating at a range of distances from predicted CPE/PIC positions. We note that a fraction of promoter windows has TSSs positions suggesting that the responsible PICs for those TSSs assemble at positions upstream or downstream from locations identified by Rhee and Pugh.

Given the distinct and polar alterations of TSS distribution at model genes by Pol II fast or Pol II slow mutants, we asked if attributes of RNA 5′ end distributions within promoter windows could also distinguish mutant classes. To do this, we examined two attributes of TSS usage: the change in position of the median TSS usage in the promoter window from WT (TSS “shift”), and the change in the width between positions encompassing 80% of the TSS usage distribution (from 10% to 90%, the change (Δ) in TSS “spread”, illustrated in **Figure 1A**). TSS shifts for individual promoters found in each mutant are displayed in a heat map that clusters by mutant profile and promoter profile (**Figure 1E**). Mutant TSS shift profiles in libraries compiled from all replicates separated strains into two major classes consistent with slow and fast Pol II mutants. Principle component analysis (PCA) of TSS shifts (**Supplemental Figure 1C**), total promoter reads (“Expression”, **Supplemental Figure 1D**), or Δ TSS spread (**Supplemental Figure 1E**) distinguish between two major classes of mutant for all individual biological replicates, corresponding to Pol II slow and fast classes. Pol II and GTF mutants showed widespread directional shifting of TSSs across nearly all promoters, with individual mutants generally shifting TSSs for most promoters either upstream (Pol II fast mutants) or downstream (Pol II slow mutants)(**Figure 1E, Supplemental Figure 1F**). Pol II GOF and *tfg2*Δ*146-180* strains exhibited primarily upstream shifts in TSS distributions within promoter windows, while Pol II LOF and *sua7-58A5* exhibited primarily downstream shifts. TSS shifts are consistent with previously observed shifts at individual promoters, such as *ADH1*, suggesting that promoter scanning is operating across all yeast promoter classes. Our analyses recapitulate a relationship between expression and TSS spread similar to that recently described for promoters from yeast, mouse, and human [87, 88]. Highly expressed promoters tend to be more focused than lowlier expressed (**Supplemental Figure 1G**). **Supplemental Figure 1H** illustrates browser tracks for the example *TUB2* promoter illustrating reproducibility at the level of individual libraries.

We examined changes in TSS distribution relative to promoter class and Pol II mutant strength to determine how each relates to magnitude of TSS changes. To visualize changes, we separated promoters using classification by Taf1-enrichment or depletion as done previously. While recent work indicates that TFIID (containing Taf1) functions at all yeast promoters [38, 40], differential recruitment of Taf1 correlates with promoter nucleosome organization, underlying DNA sequence composition, and DNA element enrichment (TATA *etc.*) [14, 18, 31, 32, 35], suggesting this metric is a useful proxy for promoter class. **Figure 2A** shows example heat maps of the difference of normalized TSS distributions between WT and a Pol II fast or a Pol II slow mutant. The stereotypical patterns of polar changes to TSS distributions where distribution of TSSs shift upstream (increases upstream and decreases downstream, such as *rpb1* E1103G), or shift downstream (increases downstream and decreases upstream, such as *rpb1* H1085Y), are observed across essentially all promoters, and for all mutants examined including GTF mutants (**Supplemental Figure 2**). By determining the shift in median TSS position in promoter windows, we can see that mutants exhibit different strengths of effects on TSS distributions (**Figure 2B**). A double mutant between *tfg2*Δ*146-180* and *rpb1* E1103G shows enhancement of TSS defects across promoter classes (**Figure 2B, 2C**), similarly to what has been observed for defects *ADH1* [79]. Counts of promoters with upstream or downstream shifts or statistical analyses for significant upstream or downstream shifts at the level of individual promoters demonstrate large directional biases in the effects of essentially all mutants (**Supplemental Figure 3**). Examination of average TSS shift and measured in vitro elongation rate for Pol II mutants shows a correlation between the strength of in vivo TSS selection defect and in vitro Pol II elongation rate [80, 81] (**Figure 2D**). These results are consistent with TSS selection being directly sensitive to Pol II catalytic activity as was suggested by our earlier work [79, 81].

**Figure 2.**
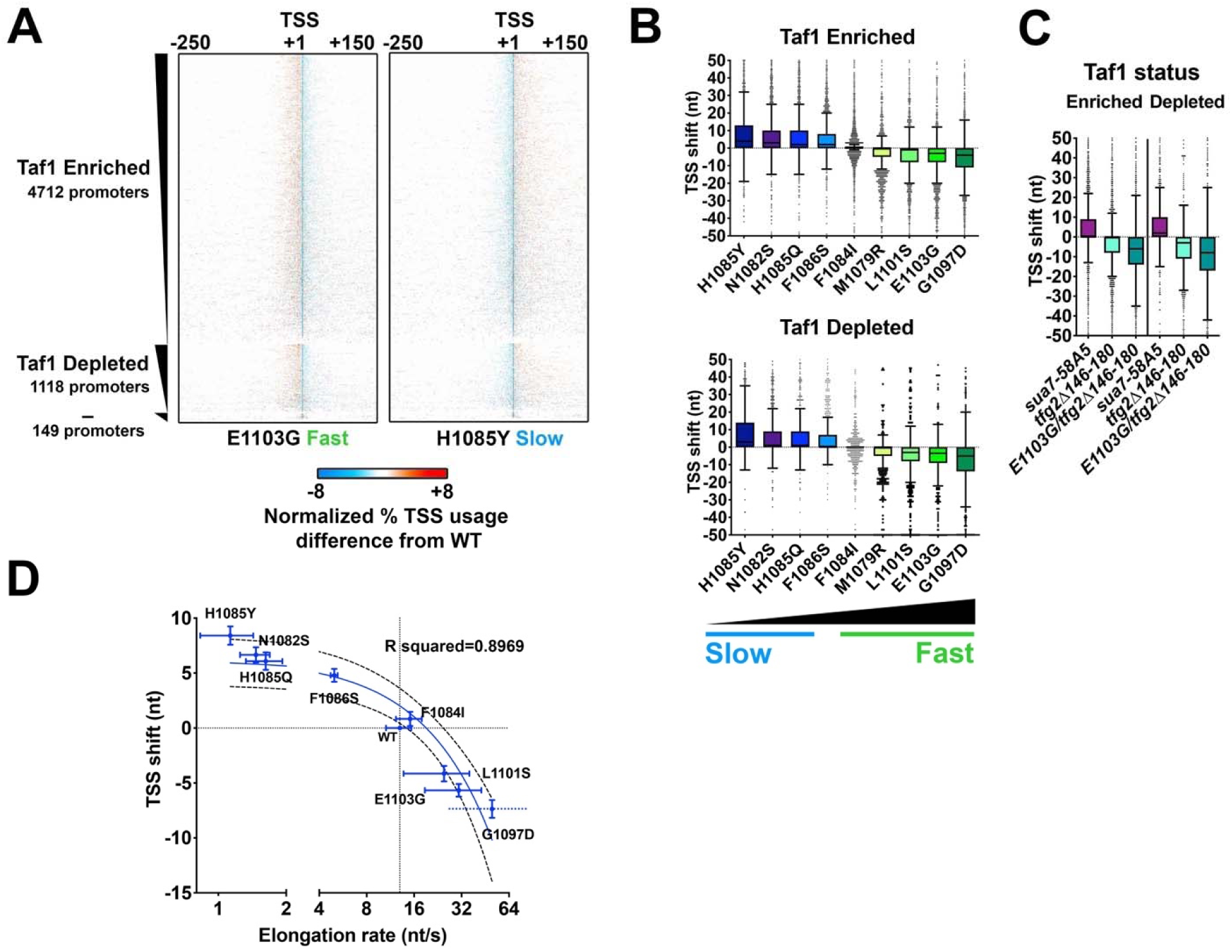
Pol II and GTF mutants confer polar shifts in TSS-usage across all promoter classes in *S. cerevisiae*. **A.** Heat maps show relative TSS distribution changes in a fast (*rpb1* E1103G) or a slow (*rpb1* H1085Y) Pol II mutant relative to WT. 401 nt promoter windows were anchored on measured median TSS position in the WT strain with TSS distributions for WT or mutant strains normalized to 100% for each promoter (heat map row). Differences in distribution between WT and mutant TSS usage were determined by subtracting the normalized WT distribution from normalized mutant distributions. Promoters are separated into those classified as Taf1-enriched (Taf1 Enriched), Taf1-depleted (Taf1 Depleted), or neither (–), and rank-ordered on the *y*-axis based on total reads in WT (from high to low). Gain in relative mutant TSS usage is positive (orange) while loss in relative mutant usage (cyan) is negative. **B.** Polar shifts in TSS usage are apparent for examined *rpb1* mutants (except *rpb1* F1084I) across promoter classes. All box plots are Tukey plots unless otherwise noted (see Methods). Promoters examined are n=3494 (>200 reads total expression in WT). Pol II mutants are rank ordered by relative strength of in vitro elongation defect (slow to fast) and colored from blue (slow) to green (fast) in similar fashion to allow visual comparison of same mutants between promoter classes. All median TSS shift values for mutants are statistically distinguished from zero at p<0.0001 (Wilcoxon Signed Rank test), except F1084I Taf1 Enriched (p=0.0021) or F1084I Taf1 Depleted (not significant) **C.** Polar shifts in TSS usage are apparent for examined GTF mutants and an *rpb1 tfg2* double mutant shows exacerbated TSS shifts relative to the single mutants (compare C to B). Promoters examined are as in (B). All median TSS shift values for mutants are statistically distinguished from zero at p<0.0001 (Wilcoxon Signed Rank test)**. D.** Average TSS shifts in Pol II *rpb1* mutants correlate with their measured in vitro elongation rates. Error bars on TSS shifts and elongation rates are bounds of the 95% confidence intervals of the means. Elongation rates are from [81, 83]. Mutants slower than WT in vitro exhibit downstream shifts in TSS distributions while mutants faster than WT in vitro exhibit upstream shifts in TSS distributions. TSS shift strength correlating with the strengths of their in vitro elongation rate defects and their in vivo growth rate defects. Linear regression line is shown along with the 95% confidence interval of the fit (dashed lines). R squared=0.8969. Note log_2_ scale on *x*-axis. Break in *x*-axis is to allow Pol II slow mutants to be better visualized. Promoters examined are as in (B,C).

**Figure 3.**
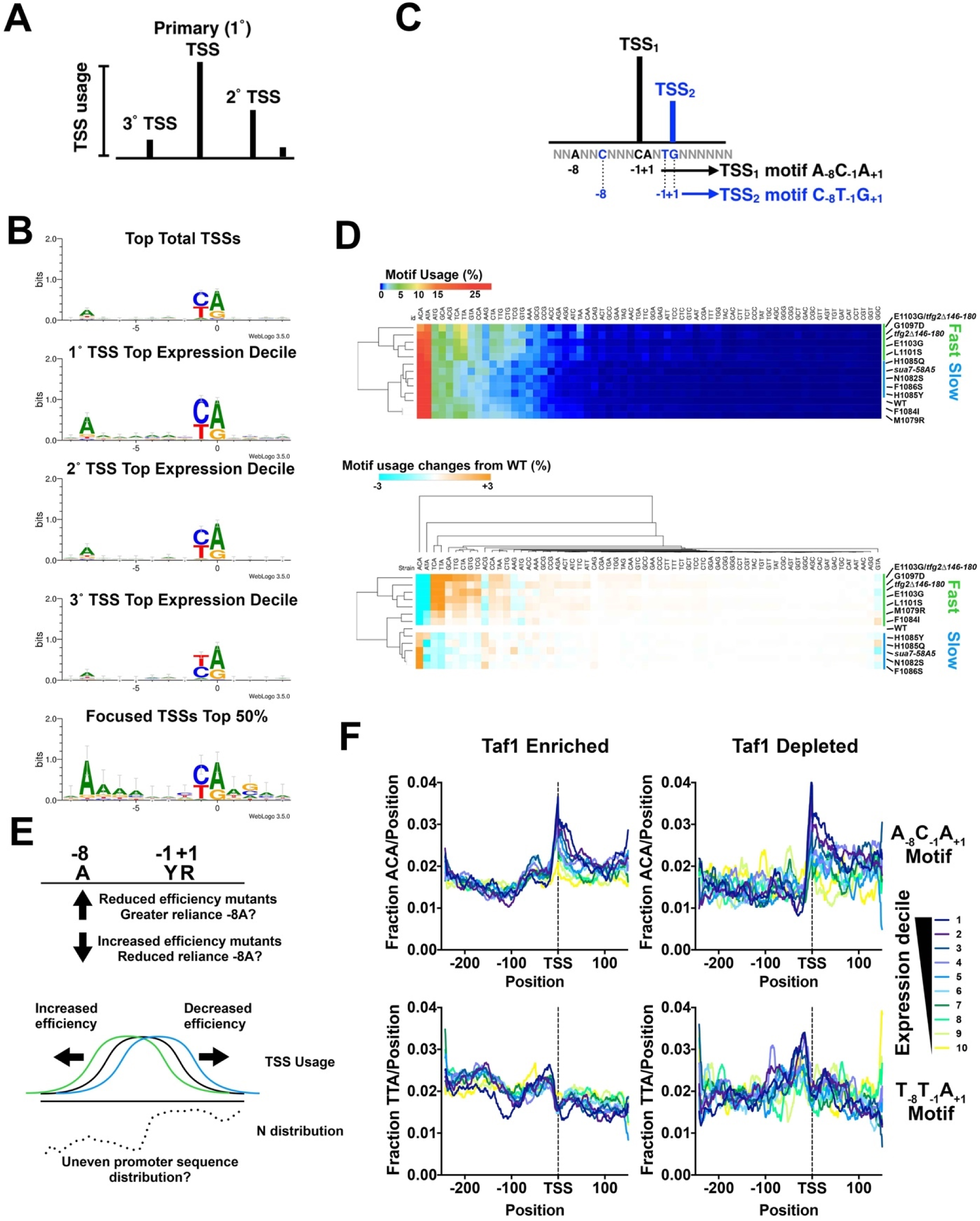
TSS motif usage and alterations in TSS-usage affecting mutants. **A.** Schematic of TSS distribution at an individual promoter defining primary TSS as most used followed by secondary and tertiary *etc* based on usage. **B.** Preferred Y_-1_R_+1_ motif usage observed in our TSS data as has been observed previously. *S. cerevisiae* selective enrichment for A at -8 is apparent at the most highly used starts in promoters with higher expression (compare primary/top (1°) TSSs with secondary (2°) or tertiary TSSs from promoters within the top decile of expression). Promoters exhibiting very narrow TSS spreads (focused) show additional minor enrichments for bases near the TSS. **C.** Schematic indicating how each TSS can be separated into one of 64 groups based on identity of nucleotides at positions -8, -1, and +1 relative to the TSS (at +1). **D.** Overall TSS motif usage in WT and TSS-usage affecting mutants. TSSs were separated by N_-8_N_-1_N_+1_ identity (64 motifs) as the vast majority of TSS reads derive from N_-8_Y_-_ 1R_+1_ sequences. This means each of 64 motifs encompasses TSSs for N_-7_N_-6_N_+5_N_-4_N_-3_N_-2_ sequences. (Top) Percent motif usage determined for individual strains and displayed in heat map hierarchically clustered on *y*-axis to group strains with similar motif usage distribution. (Bottom) Difference heat map illustrating relative changes in N_-8_Y_-1_R_+1_ motif usage in heat map hierarchically clustered on *y*-axis to group strains with similar motif usage difference distribution. **E.** Alteration in motif usage and apparent changes to reliance on an A_-8_ could arise from a number of possibilities. Alterations in TSS efficiencies in mutants could result in upstream or downstream shifts in TSS distribution if mutants have decreased or increased reliance, respectively, on a particular motif. Conversely, alteration in initiation efficiency in general (increase or decrease) could alter TSS motif usage if TSS motifs are unevenly distributed across yeast promoters (example distribution for hypothetical motif *N*). **F.** TSS motifs are unevenly distributed across yeast promoters and differentially enriched correlating with steady state promoter expression levels. (Top) the apparent highest used A_-8_Y_-1_R_+1_ motif (A_-8_C_-1_A_+1_) and (bottom) the less preferred T_-8_T_-1_A_+1_ motif were compared for Taf1 Enriched or Taf1 Depleted promoters for promoters separated into overall expression decile (Decile 1 contains highest expressed promoters, Decile 10 the lowest).

### Altered TSS motif usage in TSS-shifting mutants

To understand the basis of directional TSS shifting in Pol II mutants, we asked how changes to TSS selection relate to potential sequence specificity of initiation (**Figure 3**). Earlier studies of TSS selection defects in yeast suggested that mutants might have altered sequence preferences in the PIC [41]. Our identified TSSs reflect what has been observed previously for Pol II initiation preferences, *i.e.* the simplest TSS motif is Y_-1_R_+1_ as in most eukaryotes, with the previously observed budding yeast-specific preference for A_-8_ at strongest TSSs [43](**Figure 3B**). Preference for Y_-1_R_+1_ is common across RNA polymerases and likely reflects the stacking of an initiating purine (R, A/G) triphosphate onto a purine at the -1 position on the template strand (reflected as pyrimidine (Y, C/T) on the transcribed strand)[51]. Within the most strongly expressed promoters, preference for A_-8_ is greatest for the primary TSS, and is reduced from secondarily to tertiarily-preferred TSSs, even though these sites also support substantial amounts of initiation. Examination of the most focused, expressed promoters – promoters that contain the majority of their TSSs in a narrow window – reveals potential preferences at additional positions. We analyzed TSS usage within promoter windows by dividing all TSSs into 64 motifs based on identities of the -8, -1, and +1 positions (**Figure 3C**). We asked if Pol II or GTF mutants altered apparent preferences among these 64 motifs. Based on aggregate usage of sequences across our promoter set, we found that the top used motifs were generally A_-8_Y_-1_ R_+1_, with the next preferred motifs found among B_-8_(not A)Y_-1_R_+1_ (**Figure 3D**). Pol II and GTF mutants have clear effects on motif usage distribution concerning the -8A position. Upstream TSS shifting mutants (Pol II GOF and *tfg2*Δ*146-180*) show a decreased preference for A_-8_Y_-1_R_+1_ motifs concomitant with a gain in relative usage of B_-8_Y_-1_R_+1_ motifs, while downstream TSS shifting mutants (Pol II LOF and *sua7-58A5*) have the converse effect, though primarily increases in A_-8_C_-1_A_+1_ and A_-8_C_-1_G_+1_. Total TSS usage might be affected by strong effects at a subset of highly expressed promoters, therefore we also examined motif preference on a promoter by promoter basis (**Supplemental Figure 4A, B**). *rpb1* E1103G TSS preferences illustrate that the reduction in preference for A_-8_Y_-1_R_+1_ motifs is observed across yeast promoters (**Supplemental Figure 4A**) while H1085Y shows the converse (**Supplemental Figure 4B**).

**Figure 4.**
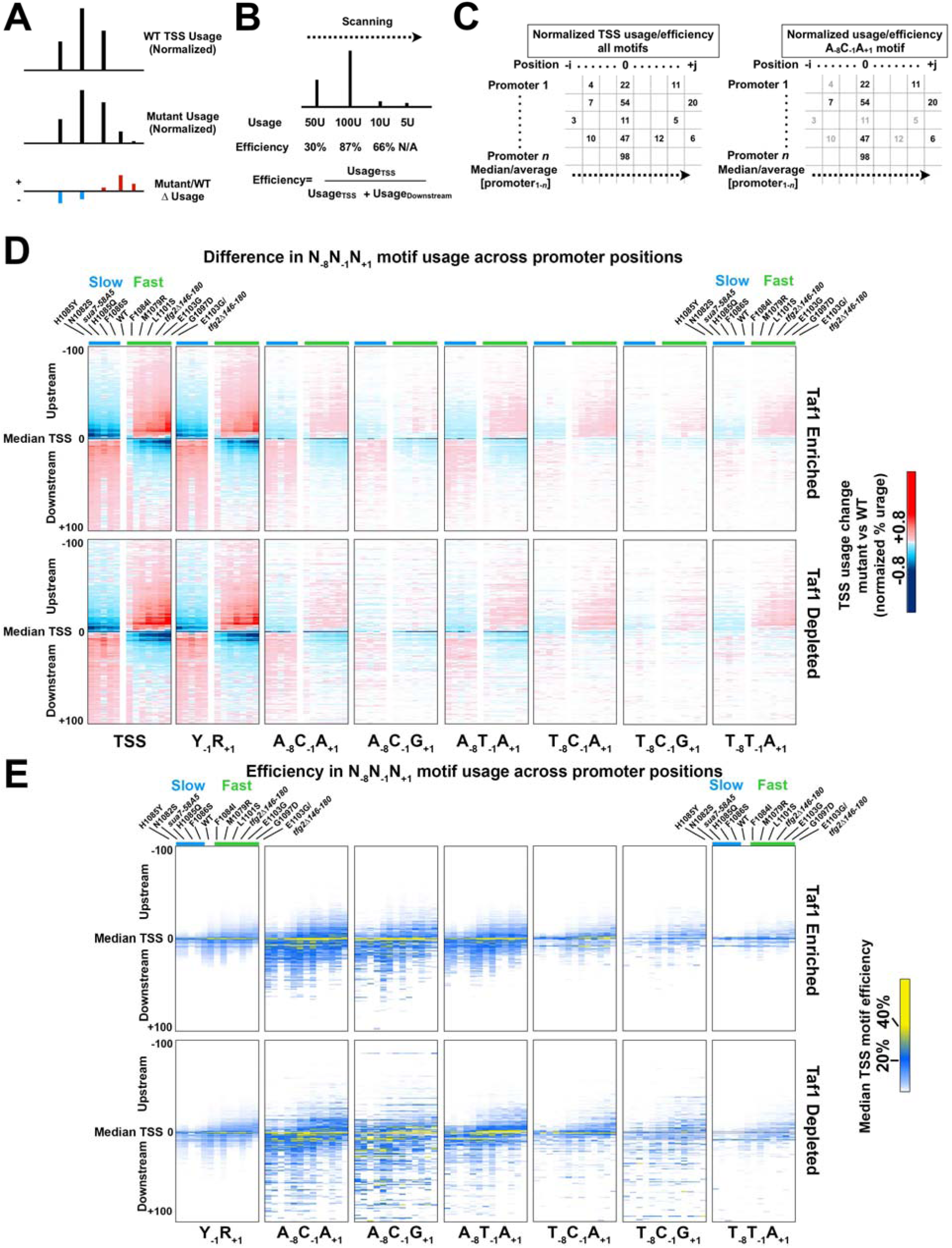
TSS-usage mutants alter TSS usage efficiencies across TSS motifs consistent with promoter scanning initiation at all promoters. **A.** Schematic indicating how normalized difference heat maps are generated (for visual purposes, differences scaled in this schematic to 1.5×). **B.** In a directional scanning mechanism, TSS efficiency is defined as the usage at a given TSS divided by that usage and all downstream usage. This allows strength of TSS to be compared instead of absolute usage, which is determined by “first come, first served” priority effects as the probability of initiation reaches a limit of one. **C.** Schematic illustrating that TSS usages/efficiencies across all promoters and positions form a matrix, and each of 64 motif TSSs represent only a subset of these values (for example A_-8_C_-1_A_+1_). Comparison of median or average values for usage/efficiency for each N_-8_N_-1_N_+1_ motif TSS subset across promoters at each promoter position allows for partial control of sequence and position variables in comparing how initiation mutants affect TSS usage. **D.** Altered usage across TSS motifs in TSS-usage affecting mutants. Heat maps show difference in aggregate usage normalized to promoter number for different N_-8_Y_-1_R_+1_ TSS motifs. Strains are ordered on the *x*-axis from left-to-right from strongest downstream shifter to strongest upstream shifter, with class of Pol II mutant (fast or slow) indicated by green and blue color bars, respectively. Promoter positions from -100 (upstream) to +100 (downstream) flanking the median TSS position in WT are shown. Regardless or promoter class, TSS-usage affecting mutants cause polar effects on distribution of TSS usage when examining motifs separately. **E.** Motif efficiency was calculated as in (B) for a subset of N_-8_Y_-1_R_+1_ TSS motifs across promoters at each promoter position for all mutants. Heat maps are ordered as in (D). Downstream shifting mutants in (D) generally *reduce* TSS usage efficiencies across promoter positions. Upstream shifting mutants in (D) generally *shift* TSS efficiencies upstream.

Different models might explain why initiation mutants alter apparent TSS sequence selectivity, and in doing so lead to polar changes to TSS distribution or vice versa (**Figure 3E**). First, relaxation of a reliance on A_-8_ would allow, on average, earlier initiation in a scanning window because non-A_-8_ sites would be encountered by the PIC at higher frequency, whereas increased reliance on A_-8_ would have the opposite effect. Alternatively, altered Pol II catalytic activity or GTF function may broadly affect initiation efficiency across all sites, which allows at least two predictions. First, an *apparent* change in TSS selectivity could result from a corresponding uneven distribution in TSS motifs within promoter regions. It has already been observed that yeast promoter classes sequence distributions deviate from random across promoters. Second, the enrichment of A_-8_Y_-1_R_+1_ TSSs used and the ability of the -8A to also function as a TSS when it is part of a YR element is strong enough that it likely underlies the prevalence for yeast TSS to be 8 nt apart [45]. Only a subset of -8As will themselves be embeded in Y_-1_R_+1_ or A_-8_Y_-1_R_+1_ elements, therefore any increase in TSS efficiencies across all sequences will be predicted to shift preference from A_-8_Y_-1_R_+1_ to B_-8_Y_-1_R_+1_. Here, we examined sequence distributions for individual nucleotides and for select A_-8_Y_-1_R_+1_ motifs relative to median TSS position for yeast promoters (**Figure 3F, Supplemental Figure 4C**). As noted previously, yeast promoter classes differ based on their distributions of A/T [35, 89]. In Wu and Li, promoters were classified based on their nucleosome structure. Our classification based on Taf1-enrichment similarly divides yeast promoters with Taf1-depleted promoters highly enriched for T and depleted for A on the top DNA strand (**Supplemental Figure 4C**). Furthermore, the extent of depletion or enrichment correlates with promoter expression level in vivo, fitting with prediction based on reporter promoter analyses [90]. Enrichment or depletion of individual nucleotides would also be expected to potentially alter distributions of N_-8_Y_-1_R_+1_ TSS motifs. Therefore, we extended our analyses to N_-8_Y_-1_R_+1_ motifs (**Figure 3F**). We find that A_-8_C_-1_A_+1_, the apparent most-preferred TSS motif for Pol II in yeast, is markedly enriched at the median TSS and downstream positions with a sharp drop off upstream, with enrichment also showing correlation with apparent promoter expression level. A less preferred motif, T_-8_T_-1_A_+1_, shows a distinct enrichment pattern (enriched upstream of median TSS, depleted downstream). This biased distribution in promoter sequence for TSS sequence motifs makes it difficult to determine whether apparent altered sequence specificity is a cause or consequence of altered TSS distributions.

### TSS motif efficiency and usage altered across a number of TSS motifs

To examine further, we looked at the overall shapes of TSS distributions to determine if mutants alter the shapes of TSS distributions or merely shifted them (**Figure 4**). To do this, we examined overall usage and usage for particular TSS motifs at promoters but also efficiencies of TSS usage for individual TSS motifs (**Figure 4A, 4B**). Efficiency is determined as the ratio of observed reads for a particular TSS to the sum of those reads and all downstream reads, as defined by Kuehner and Brow [61] (**Figure 4B**). A scanning mechanism predicts first come-first served behavior in observed TSS usage dependent on innate efficiency of a given TSS (**Figure 4B**). Scanning from upstream to downstream will create greater apparent usage for upstream TSSs relative to a downstream TSS, even if they are equally strong in promoting initiation. If Pol II mutants primarily affect initiation *efficiency* across TSSs, we have specific expectations for how TSS distributions will be affected. For example, if slow Pol II alleles decrease initiation efficiency across sequences, we predict that usage distribution will be flatter than WT. This “flatness” will appear as a downstream shift in usage, and result in the median observed TSS efficiency being lower than WT over all promoter positions except for the very downstream tail of usage. This would reflect a spreading out of the usage distribution to downstream positions as fewer Pol II molecules would initiate at upstream positions, and more Pol II would continue to scan to downstream relative to WT. Conversely, if fast Pol II alleles increase initiation efficiency across sequences, we would predict that both TSS usage and median efficiency to increase for upstream promoter positions but return to baseline efficiency sooner than WT.

To partially account for innate sequence differences among TSS motifs, we examined TSS usage and efficiency across promoters for specific N_-8_Y_-1_R_+1_ motifs (**Figure 4C, Supplemental Figure 5**). Usage is defined as the reads found in particular TSS relative to the total reads for that promoter, whereas efficiency is an estimate of the strength of a TSS, assuming a polar scanning process as illustrated in **Figure 4B**. Extending this motif analysis to a range of N_-8_Y_-_ 1R_+1_ motifs used at different levels (**Figure 4D, 4E, Supplemental Figure 5A-D**) we observe that upstream-shifting mutants shift usage upstream for all examined motifs (**Figure 4D**) while downstream-shifting mutants have the opposite effects on motif usage for all examined motifs. In contrast, when examining N_-8_Y_-1_R_+1_ motif efficiencies across promoter positions, downstream-shifting mutants tend to reduce efficiencies across promoter positions while upstream-shifting mutants shift TSS efficiencies upstream (**Figure 4E**). These analyses are consistent with upstream-shifting mutants exhibiting increased efficiency across TSS motifs and promoter positions, which shifts both the usage and observed efficiency distributions to upstream positions, while downstream mutants reduce the efficiency curve and essentially flatten the usage distributions, as would be expected from reduced initiation efficiency across promoter positions. Analysis indicates broad statistical significance for TSS usage and efficiency effects for examined *rpb1* H1085Y and E1103G mutants across promoter positions and TSS motifs (**Supplemental Figure 5C, 5D**).

**Figure 5.**
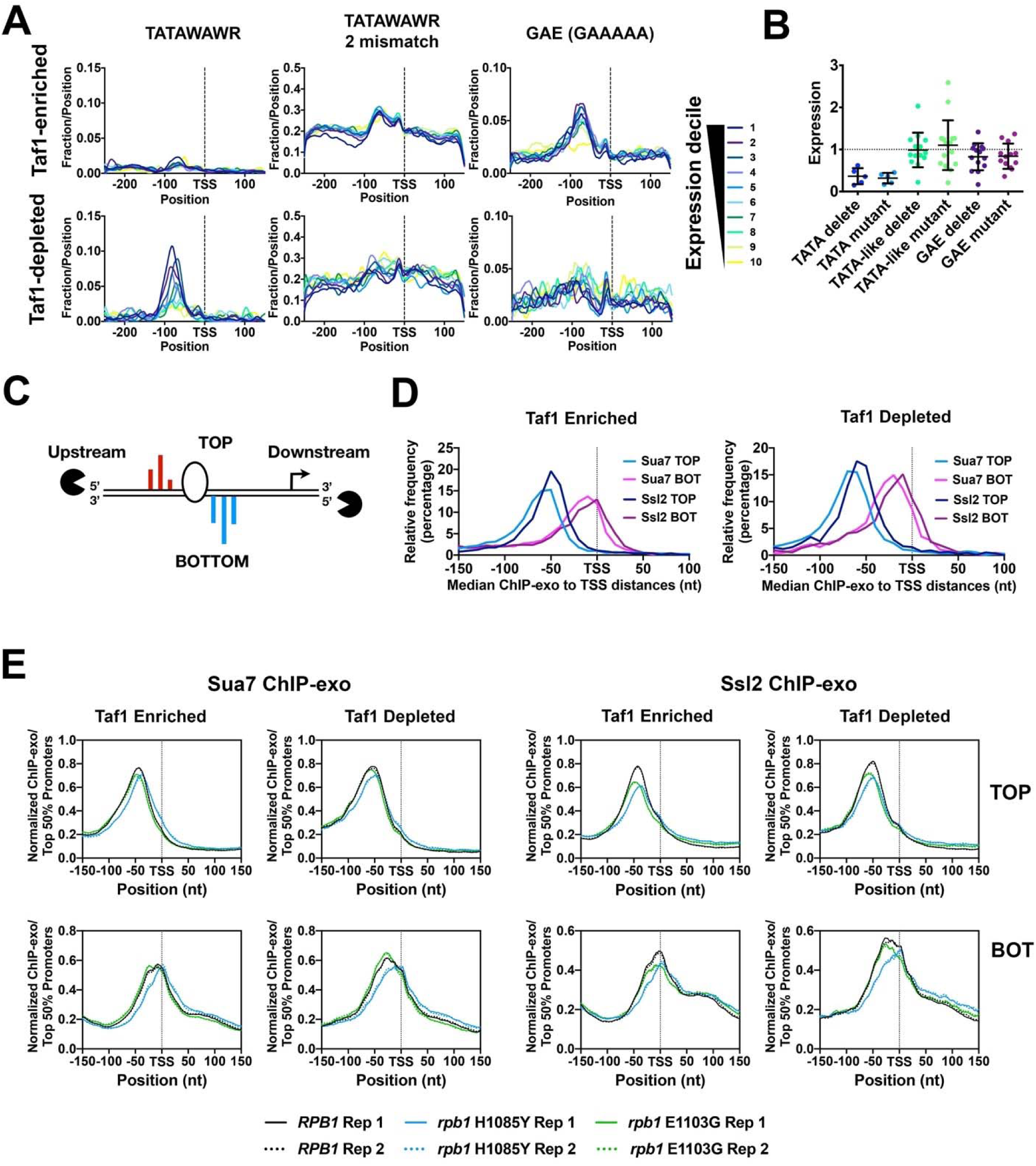
Attributes of core promoter classes and PIC positioning in TSS-usage affecting mutants. **A.** Enrichment by expression decile in WT of putative core promoter elements in Taf1 Enriched and Taf1 Depleted promoters. TATA consensus (TATAWAWR, W=A/T, R=A/G) is enriched in Taf1 Depleted promoters while the GA-rich element (GAAAAA) is enriched in Taf1 Enriched promoters. Yeast promoters are relatively AT-rich so there is a high probability of “TATA-like” elements differing from the TATA consensus by two mismatches. **B.** Tested GAE or TATA-like elements do not greatly contribute to expression from promoters where tested. Expression by Northern blotting for promoters or classes of promoter mutant fused to a reporter gene. Promoter mutants were normalized to the respective WT version of each promoter. “Delete” mutants represent deletions of particular element types. “Mutant” elements represent elements where base composition has been altered. **C.** GTF positioning by promoter classes determined by ChIP-exo for Sua7 (TFIIB) or Ssl2 (TFIIH). For each promoter, the median position of ChIP-exo reads on the top (TOP) or bottom (BOT) DNA strand was used to estimate GTF positioning. TOP and BOT strands are defined relative to promoter orientation in the genome and have the same upstream and downstream as a promoter. **D.** Left graph shows histograms of GTF signal median positions for ChIP-exo read distributions at Taf1 Enriched promoters while right graph shows histograms of GTF signal median positions for ChIP-exo read distributions at Taf1 Depleted promoters. **E.** Pol II mutant effects on GTF positioning as detected by ChIP-exo for Sua7 (TFIIB) or Ssl2 (TFIIH). Aggregate ChIP-exo signal for Taf1-enriched or depleted promoters on top (TOP) or bottom (BOT) DNA strands in WT, *rpb1* H1085Y, or *rpb1* E1103G. Curves on graph indicate 2^nd^ order polynomial (10 neighbor) smoothing of promoter-normalized ChIP-exo reads averaged for the top 50% of promoters determined by ChIP-exo reads in WT cells. Biological replicate data are shown for each strain and replicates are essentially superimposable.

### Analysis of promoter architecture to understand location of PIC assembly and estimate scanning distances for yeast promoters

High-resolution TSS data allow us to evaluate promoter features and their potential relationships to observed median TSS positions instead of using annotated TSS (one per gene and not necessarily accurate) from the Saccharomyces Genome Database. For example, in a scanning mechanism, TSSs may have evolved at different distances from the point of scanning initiation. This would mean that different promoters may have different scanning distances, which could result in differential sensitivity to perturbation to initiation. As has previously been determined, a minority of yeast promoters contain consensus TATA elements (TATAWAWR) and these are enriched in Taf1-depleted promoters (illustrated in **Figure 5A**) within ∼50-100 basepairs upstream of TSS clusters. Furthermore, TATA enrichment tracks with apparent expression level determined by total RNA 5′ reads within promoter windows. For this class of promoter, a consensus TATA element seems the likely anchor location for PIC assembly and the determinant for the beginning of the scanning window. However, TATAWAWR elements are not enriched in Taf1-enriched promoters. On the basis of finding TATA-like elements within ChIP-exo signal for GTFs along with a putative stereotypical pattern to the ChIP-exo signal, it has been proposed by Rhee and Pugh that promoters lacking consensus TATA elements can use TATA-like elements (TATAWAWR with one or two mismatches) for function analogous to a TATA element [14]. Therefore, such elements might potentially serve as core promoter elements anchoring PIC formation and determining the promoter scanning window for these promoters. Evidence for the function of such TATA-like elements is sparse. In vitro experiments suggested that a TBP footprint is positioned over potential TATA-like element in *RPS5* promoter, but the element itself is not required for this footprint [91]. In contrast, more recent results have suggested modest requirement for TATA-like elements at three promoters (∼2-fold) in an in vitro transcription system [92]. Examination of the prevalence of elements with two mismatches from TATA consensus TATAWAWR within relatively AT-rich yeast promoter regions suggests that there is a high probability of finding a TATA-like element for any promoter (**Figure 5A**). Taf1-enriched promoters show enrichment for an alternate sequence motif, a G-capped A tract (sequence GAAAAA), also called the GA-element (GAE) [35, 36]. This positioning of GAEs approximately 50-100 bp upstream of TSSs is reminiscent of TATA positioning (**Figure 5A**), and the GAE has been proposed to function as a core promoter element at non-TATA promoters [36]. Other studies describe the relationship of this element to nucleosome positioning and suggest that these elements may function directionally in nucleosome remodeling at NDR promoters as asymmetrically distributed poly dA/dT elements [93, 94]. To understand if these potential elements function in gene expression, which would be predicted if they served as potential PIC assembly locations, we cloned a number of candidate promoters upstream of a *HIS3* reporter and deleted or mutated identified TATA, TATA-like, or GAE elements and examined effects on expression by Northern blotting (**Figure 5B, Supplemental Figure 6**). As expected, in general, identified consensus TATAs positioned upstream of TSSs were important for normal expression of the *HIS3* reporter. In contrast, neither TATA-like or GAE elements in general had strong effects on expression, though some individual mutations affected expression to the same extent as mutation of TATA elements in the control promoter set. We conclude that GAE or TATA-like elements do not generally function similarly to consensus TATAs for promoter expression.

**Figure 6.**
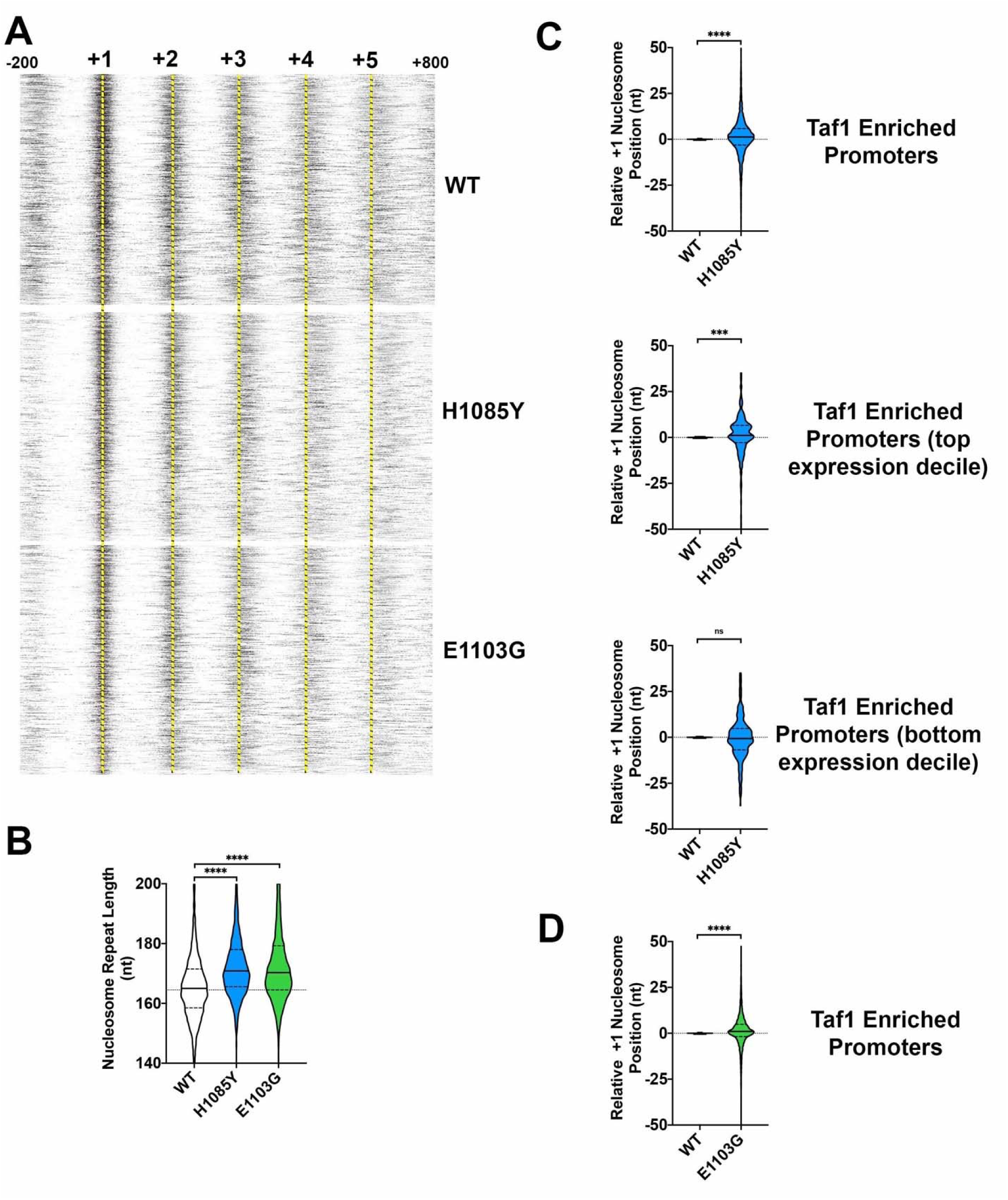
Effects of slow and fast Pol II mutants on nucleosome positioning. **A.** Average nucleosome midpoints per promoter from MNAse-seq for WT, *rpb1* H1085Y, and *rpb1* E1103G were mapped for Taf1 Enriched promoters anchored on experimentally determined +1 nucleosome positions at +1 (−200 to +800 positions shown). Both *rpb1* H1085Y and *rpb1* E1103G shift genic nucleosomes downstream relative to WT. WT average nucleosome positions determined by autocorrelation analysis of WT nucleosome midpoints. Data are from two (WT), seven (*rpb1* H1085Y), or eight (*rpb1* E1103G) independent biological replicates. Yellow dashed lines allow comparison of WT nucleosome positions with *rpb1* mutants. **B.** Nucleosome repeat lengths determined by autocorrelation analysis on the independent replicates noted in (A). Both *rpb1* H1085Y and *rpb1* E1103G nucleosome repeat lengths are significantly different from WT (Wilcoxon Matched-Pairs Signed Rank test, two tailed, p<0.0001). **C.** +1 nucleosome positioning in *rpb1* H1085Y is subtly altered from WT for Taf1 Enriched promoters. Top violin plot shows the distribution of individual +1 nucleosomes for *rpb1* H1085Y biological replicates (n=7) relative to the position determined from the WT average (n=2) for Taf1 Enriched promoters (n=4161). +1 nucleosome positions are significantly different (Wilcoxon Matched-Pairs Signed Rank test, two tailed, p<0.0001). Middle violin plot as in top but for Taf1 Enriched promoters in the top expression decile (n=321). +1 nucleosome positions are significantly different (Wilcoxon Matched-Pairs Signed Rank test, two tailed, p=0.0004). Bottom violin plot as in middle but for Taf1 Enriched promoters in the lowest expression decile (n=376). +1 nucleosome positions are not significantly different (test as above). **D.** +1 nucleosome positioning in *rpb1* E1103G is subtly altered from WT for Taf1 Enriched promoters. Violin plot shows the distribution of individual +1 nucleosomes for *rpb1* E1103G biological replicates (n=8) relative to the position determined from the WT average (n=2) for Taf1 Enriched promoters (n=4161). +1 nucleosome positions are significantly different (Wilcoxon Matched-Pairs Signed Rank test, two tailed, p<0.0001).

### TSS-shifting initiation mutants alter PIC-component positioning consistent with promoter scanning model

Given results above suggesting that TATA-like or GAE elements may not generally function as core promoter elements and therefore may lack value as potential PIC landmarks, we performed ChIP-exo for GTFs TFIIB (Sua7) and TFIIH (Ssl2) to directly examine PIC component localization in WT, *rpb1* H1085Y, and *rpb1* E1103G cells (**Figure 5C**). Element-agnostic analyses of ChIP-exo [95] for Sua7 and Ssl2 was performed in duplicate for all strains. ChIP-exo v5.0 signal was highly reproducible (**Supplemental Figure 7A-B**). We reasoned that ChIP-exo would allow us to determine where the PIC localizes for all promoter classes and, moreover, how PIC localization may be altered by Pol II mutants that alter TSS utilization. As discussed above, previous work anchored ChIP-exo signal for PIC components over TATA or TATA-like sequences and identified a stereotypical overall pattern for crosslinks relative to these anchor positions. These crosslink patterns were interpreted as relating to potential structure of the PIC open complex [14]. Subsequent work has identified that crosslinking in ChIP-exo can have some sequence bias [96] and this sequence bias may reflect partially the stereotypical crosslinking patterns observed around TATA/TATA-like sequences. Because the PIC must access TSSs downstream from the site of assembly, it is likely that observed ChIP-exo signal reflects the occupancies of PIC components across promoters and not only the site(s) of assembly. Using TATA-like sequences as anchors, Taf1-enriched promoters were found to have PIC components on average closer to TSSs than they were for Taf1-depleted promoters [14]. Here, we used our high resolution TSS mapping data coupled with determination of median position of ChIP-exo signal for Ssl2 or Sua7 within promoter windows to examine distance between putative PIC position and initiation zone as reflected by observed median TSSs (**Figure 5C-E**). **Figure 5C** illustrates basic concepts of ChIP-exo in that the exonuclease approaches crosslinked promoter complexes from the upstream direction on the top DNA strand of a promoter and from the downstream direction on the bottom strand. Top and bottom strands are organized with the same upstream and downstream directions as they indicate the two DNA strands of a directional promoter region. Using median ChIP-exo signal within promoter windows for Ssl2 or Sua7 on top or bottom promoter strands (TOP or BOT), we find that this simple metric behaves as predicted for PIC component signal (**Figure 5D**). **Figure 5D** shows the histogram for individual promoter median ChIP-exo positions for components on the two promoter strands Sua7 signal is slightly upstream of Ssl2 signal, as expected for upstream and downstream components of the PIC, though there is considerable overlap in signal if considering TOP-BOT distance. We also confirm that on average, ChIP-exo signal for PIC components is closer to median TSS position for Taf1-enriched promoters than for Taf1-depleted promoters.

**Figure 7.**
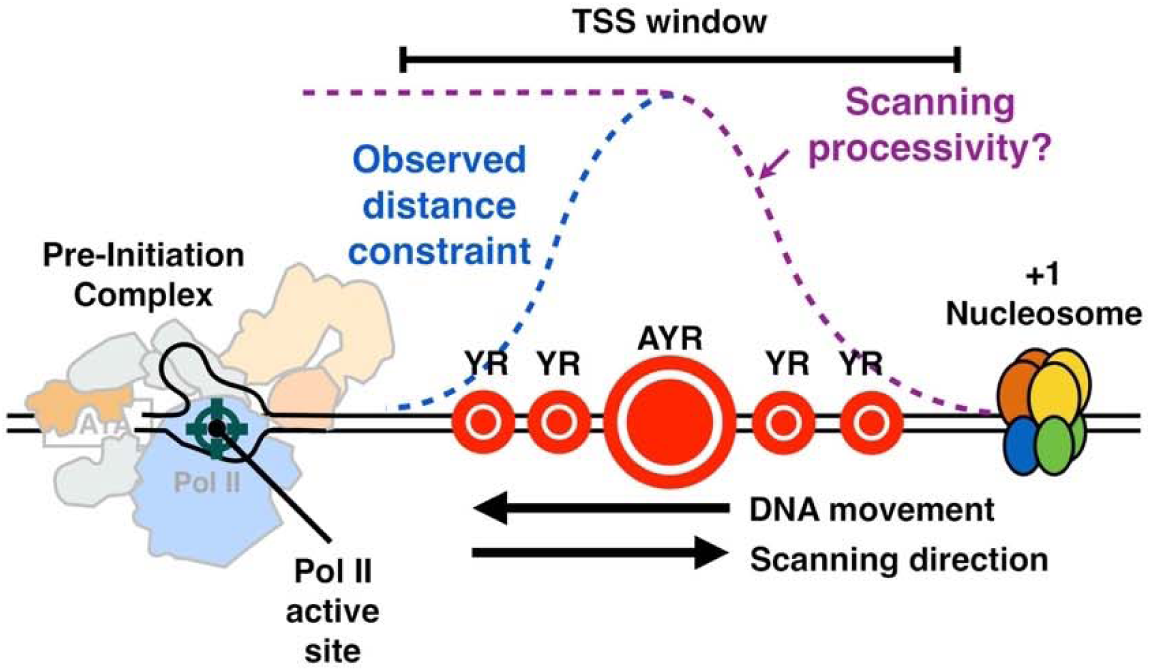
“Shooting Gallery” model for initiation by Pol II scanning in *Saccharomyces cerevisiae*. PIC assembly upstream of TSS region initiates a scanning process whereby TSSs are moved toward the PIC by DNA translocation putatively through TFIIH DNA translocase activity. Initiation probability will be determined in part by DNA sequence (the size of the indicated “targets”), Pol II catalytic activity, and the processivity of scanning, as well as constraints of TSSs being too close to the PIC. Our data are consistent with this mechanism acting at all yeast promoters and enable interpretation of how alterations to Pol II catalytic activity, TFIIF function, or TFIIB function alter initiation probability at all TSSs.

We reasoned that if ChIP-exo signal for PIC components at least partially reflects promoter scanning, *i.e.* the interaction of PIC components with downstream DNA between PIC assembly position and the zone of initiation, then Pol II mutants that alter TSS usage distribution should also alter PIC component distribution across promoters. We observed changes to the aggregate distribution of ChIP-exo signal for both Taf1-enriched and Taf1-depleted promoter classes, with effects most obvious on the downstream edge of the PIC as detected by Ssl2 signal on the bottom strand of promoter DNA, especially for *rpb1* H1085Y (**Figure 5E, Supplemental Figure 7A-C**). The shifts observed in aggregate are also observed if we look at shifts for promoter ChIP-exo medians individually (**Supplemental Figure 7A-C**). In single molecule experiments examining putative promoter scrunching in the Pol II PIC, scrunching behavior was similar regardless of whether all NTPs (to allow initiation) were present [97]. This observation suggested the possibility that putative promoter scanning driven by TFIIH-mediated scrunching might be uncoupled from initiation. In other words, that TFIIH translocation might continue independently of whether Pol II initiates or not. However, we observed altered PIC component localization in Pol II mutants predicted to directly alter initiation efficiency but not necessarily other aspects of PIC function such as TFIIH-mediated scanning (directly). Thus, there may in fact be coupling of initiation and scanning in vivo. Apparent coupling has been observed in magnetic tweezers experiments where a short unwinding event that is strictly TFIIH-dependent can be extended to a larger unwinding event by addition of NTPs, presumably reflecting Pol II transcription [98].

### Relationships of TSS-selection altering initiation mutants with promoter architectural features

TSSs evolve at certain distances from the site of PIC assembly. This means that TSSs will be found at a range of distances from sites of initial assembly and will theoretically require scanning of different distances. We asked whether presumed scanning distance correlated with promoter sensitivity to Pol II mutants for TSS shifts (**Supplemental Figure 8**). We observed at most a very modest correlation for TSS shifting extent based on where TSSs are relative to PIC location for Taf1 Enriched promoters (**Supplemental Figure 8A**). Even where correlation shows strong significance, such correlation explains only a small fraction of TSS shift relative to ChIP-exo positions. However, greater correlation between TSS shift in initiation mutants and ChIP-exo signal was observed for Taf1 Depleted promoters having consensus TATA elements (**Supplemental Figure 8B**). These latter promoters have putative PIC assembly points at greater distances from TSSs on average. Within the range of distances where most of these promoters have their TSSs, promoters with TSSs evolved at downstream positions show the greatest effects of upstream-shifting mutants on the TSS distribution (the TSS shift). Conversely, promoters with TSSs evolved at upstream positions show the greatest effects of downstream shifting mutants. These results are consistent with a facet of promoter architecture correlating with altered initiation activity, but with potential upstream and downstream limiters on this sensitivity (see Discussion for more).

The majority of yeast promoters, especially the Taf1 Enriched class, are found within an NDR and flanked by an upstream (−1) and a downstream (+1) nucleosome. Previous work showed association between ChIP-exo for GTFs and +1 nucleosomes [14], and our data illustrate this as well (discussed below). ChIP-exo for PIC components tracks with nucleosome position with some flexibility. How the PIC recognizes promoters in the absence of a TATA-box is an open question. These results are consistent with the fact that TFIID has been found to interact with nucleosomes [99] and with the possibility that the +1 nucleosome may be instructive for, or responsive to, PIC positioning. Nucleosomes have previously been proposed as barriers to Pol II promoter scanning to explain the shorter distance between PIC-component ChIP-exo footprints and TSSs at Taf1 Enriched promoters [14]. Nucleosomes can be remodeled or be moved by transcription in yeast [15, 100], likely during initiation as even for promoters with NDRs, TSSs can be found within the footprints of the +1 nucleosome. We do not observe a differential barrier for downstream shifting in Pol II or GTF mutants at Taf1-enriched promoters, which have positioned nucleosomes (**Figure 2B**), thus it remains unclear whether the +1 nucleosome can act as a barrier for Pol II scanning or TSS selection from the existing data.

To determine if altered initiation and PIC positioning of Pol II mutants, especially downstream shifting *rpb1* H1085Y, occurs in conjunction with altered +1 nucleosome positioning, we performed MNase-seq in *rpb1* H1085Y and E1103G mutants along with a WT control strain (**Figure 6, Supplemental Figures 9-10**). Determination of nucleosome positioning by MNase-seq can be sensitive to a number of variables (discussed in [101]), therefore we isolated mononucleosomal DNA from a range of digestion conditions and examined fragment length distributions in MNase-seq libraries from a number of replicates (**Supplemental Figure 9A**) to ensure we had matched digestion ranges for WT and mutant samples. Our data recapitulate the observed relationship PIC component and nucleosome positioning (**Supplemental Figure 9B, 9C**)[14]. Nucleosomes and PIC component signal are correlated but in an intermediate fashion relative relative to PIC component-median TSS correlation. We asked if +1 nucleosome midpoints were affected in aggregate, if array spacing over genes was altered, or if individual +1 nucleosomes shifted on average in Pol II mutants vs. WT. Aligning genes of Taf1 Enriched promoters by the +1 nucleosome position in WT suggests that both *rpb1* H1085Y and *rpb1* E1103G nucleosomes show significantly increased nucleosome repeat length, which becomes visually obvious at the +3, +4, and +5 positions relative to WT (**Figure 6A, 6B, Supplemental Figure 10A, 10B, 10H**). For *rpb1* H1085Y, we observed a slight but apparently significant shift for the aggregate +1 position (**Figure 6C**, top). The downstream shift in aggregate +1 position also is reflected at the individual nucleosome level across H1085Y replicates (violin plots, **Supplemental Figure 10C**). To ask if this effect on nucleosomes reflected a global defect across genes or instead correlated with transcription (whether it be initiation or elongation), we performed the same analyses on the top expression decile (**Figure 6C**, middle, **Supplemental Figure 10D,E**) and bottom expression decile Taf1 Enriched promoters (**Figure 6C**, bottom, **Supplemental Figure 10F,G**). The downstream shift was apparent in top expression decile promoters but not in bottom expression decile promoters, as would be predicted if the alteration were coupled to transcription. For *rpb1* E1103G, we observed a slight shift (∼1nt)(**Figure 6D, Supplemental Figure 10H,I**). To potentially identify subpopulations of nucleosomes, we employed a more sophisticated analysis of nucleosomes using approach of Zhou *et al* [101] (**Supplemental Figure 9B**). This approach recapitulated a similarly slight effect of H1085Y on shifting the +1 nucleosome downstream across most H1085Y datasets relative to WT.

## DISCUSSION

Budding yeast has been a powerful model for understanding key mechanisms for transcription by Pol II. An early identified difference in promoter behavior for yeast TATA-containing promoters from classically studied TATA-containing human viral promoters such as adenovirus major late led to proposals that initiation mechanisms were fundamentally different between these species [55, 102]. TSSs for yeast TATA promoters were found downstream and spread among multiple positions while TSSs for viral and cellular TATA promoters were found to be tightly positioned ∼31 nt downstream of the beginning of the element [56]. This positioning for TSSs at TATA promoters holds for many species including *S. pombe* [103] but not budding yeast. This being said, genome-wide studies of initiation indicate that the vast majority of promoters use multiple TSSs, though evolution appears to restrict TSS usage at highly expressed promoters in multiple species, including budding yeast (our work, [30, 87, 89]). How these TSSs are generated and if by conserved or disparate mechanisms is a critical unanswered question in gene expression.

We have shown here that Pol II catalytic activity, as determined by mutations deep in the active in the essential “trigger loop”, confer widespread changes in TSS distributions across the genome regardless of promoter type. Mutants in core Pol II GTFs TFIIB (*sua7* mutant) or TFIIF (*tfg2* mutant) confer defects of similar character to downstream shifting or upstream shifting Pol II alleles, respectively. The changes observed are consistent with a model (**Figure 7**) wherein TSSs are displayed to the Pol II active site directionally from upstream to downstream, with the probability of initiation controlled by the rate at which sequences are displayed (scanning rate), and by Pol II catalytic rate. This system is analogous to a “shooting gallery” where targets (TSSs) move relative to a fixed firing position (the Pol II active site)[104]. In this model, Pol II catalytic activity, the rate of target movement, *i.e.* scanning rate, and the length of DNA that can be scanned *i.e.* scanning processivity, should all contribute to initiation probability at any particular sequence. Biochemical potential of any individual sequence will additionally contribute to initiation efficiency. Our results suggest that Pol II and tested GTF mutants affect initiation efficiency across sequence motifs and that differential effects in apparent motif usage genome-wide likely result from skewed distributions of bases within yeast promoters. Our in vivo results are consistent with elegant in vitro transcription experiments showing reduction of ATP levels (substrate for initiating base or for bases called for in very early elongation) confers downstream shifts in start site usage [105]. Reduction in substrate levels in vitro, therefore, is mimicked by reduction of catalytic activity in vivo.

How template sequence contributes to initiation beyond positions close to the template pyrimidine specifying the initial purine, and how they interact with scanning, is an open question. For models employing a scanning mechanism such as the “shooting gallery”, it can be imagined that bases adjacent to the TSS affect TSS positioning to allow successful interaction with the first two NTPs, while distal bases such as the -8T on the template strand (−8A on the non-template strand) stabilize or are caught by interaction with the yeast TFIIB “reader” to hold TSSs in the active site longer during scanning [54]. Critical to this model are the structural studies just cited of Sainsbury *et al* [54] on an artificial initial transcribing complex showing direct interaction of Sua7 D69 and R64 and -8T and -7T on the template strand. There are a number of ways TFIIB may alter initiation efficiency beyond recognition of upstream DNA. TFIIB has also been proposed by Sainsbury et al to allosterically affect Pol II active site Mg^2+^ binding and RNA-DNA hybrid positioning [11, 54]. Direct analysis of Kuehner and Brow [61] found evidence for lack of effect of *sua7* R64A on efficiency of one non- -8A site, while -8A sites were affected, consistent with this residue functioning as proposed. We isolated individual motifs to examine efficiency (**Figure 4C**), and our tested *sua7-58A5* allele reduced efficiencies of both -8A and non--8A motifs alike. This allele contains a five-alanine insertion at position 58 in Sua7, likely reducing efficiency of the B-reader but possibly leaving some R64 interactions intact. Specific tests of Sua7 R64 mutants under controlled promoter conditions will directly address whether this contact confers TSS selectivity. Additionally, altered selectivity alleles of Sua7 would be predicted if interactions with the template strand were altered.

Core transcriptional machinery for Pol II initiation is highly conserved in eukaryotes leading to the general expectation that key mechanisms for initiation will be conserved. While it has long been believed that budding yeast represents a special case for initiation, this has not systematically been addressed in eukaryotes. The question of how broadly conserved are initiation mechanisms in eukaryotic gene expression is open for a number of reasons. There are examples of diverse transcription mechanisms within organisms across development, for example tissues, cells, or gene sets using TBP-related factors to replace TBP in initiation roles. For example, in zebrafish, distinct core promoter “codes” have been described for genes that are transcribed in oocytes (maternal transcription) versus those transcribed during zygotic development (zygotic transcription) [106]. The maternal code is proposed to utilize an alternate TBP for initiation, while zygotic promoters utilize TBP. Distinct core promoters are used to drive maternal and zygotic expression. For genes transcribed both maternally and zygotically, distinct TSS clusters specific to each phase of development can be quite close to one another in the genome and may have superficially similar distribution characteristics, for example promoter widths or spreads. Comparison of TSS distributions using analyses aware of distribution of possible TSSs would be a powerful tool to probe initiation mechanisms.

Another major question is how promoters without TATA-elements are specified. Organization of PIC components is relatively stereotypical within a number of species, as detected by ChIP methods for Pol II and GTFs [14, 107, 108], with the caveat that these are population-based approaches. The most common organization for promoters across examined eukaryotes is an NDR flanked by positioned nucleosomes. Such NDRs can support transcription bidirectionally, reflecting a pair of core promoters with TSSs proximal to the flanking nucleosomes [20, 21, 24-26, 109-111]. While sequence elements have been sought for these promoters, an alternate attractive possibility is that NDR promoters use nucleosome positioning to instruct PIC assembly. The association of TSSs with the edges of nucleosomes is striking across species, though in species with high levels of promoter proximal pausing, nucleosomes may be positioned downstream of the pause. Transcription itself has been linked to promoter nucleosome positioning, turnover, or exchange in yeast (for example, see [100]). Given that MNase analyses reflects bulk nucleosome populations, and depending on kinetics of initiation and the duration of chromatin states supporting initiation (expected to be relatively infrequent), the nature of initiating chromatin is unclear.

Finally, how does initiation interact with nucleosomes? In a scanning model, Pol II activity will not be expected to control the interactions with the downstream nucleosome. Instead, TFIIH bound to downstream DNA and translocating further downstream to power scanning, will be expected to be the major interaction point of the PIC and the +1 nucleosome. This model explains why downstream nucleosomes may not limit changes to scanning incurred by alterations to Pol II activity, because Pol II will be acting downstream of the TFIIH-nucleosome interaction. DNA translocation by TFIIH is expected to be competitive with the +1 nucleosome for DNA as scanning proceeds into the territory of the nucleosome. Indeed, transcription and TFIIH activity are proposed to drive H2A.Z exchange in the +1 nucleosome [100]. How TFIIH activity is controlled to either allow scanning in addition to promoter opening or be restricted to promoter opening is a major question in eukaryotic initiation. The *S. cerevisiae* CDK module of TFIIH has been implicated in restricting initiation close to the core promoter in vitro, but no evidence has emerged in vivo for this mechanism [112]. TFIIH components have long been implicated in controlling activities of the two ATPases – Ssl2 and Rad3 in yeast, XPB and XPD in humans – to enable or promote transcription or nucleotide excision repair [113-115]. These inputs may regulate activity of ATPases and their ability to be coupled to translocation activity analogous to paradigms for DNA translocase control in chromatin remodeling complexes [116].

## METHODS

### Yeast strains, plasmids, and oligonucleotides

Yeast strains used in this study were constructed as described previously [79-81]. Briefly, plasmids containing *rpo21/rpb1* mutants were introduced by transformation into a yeast strain containing a chromosomal deletion of *rpo21/rpb1* but with a wild type *RPO21/RPB1 URA3* plasmid, which was subsequently lost by plasmid shuffling. GTF mutant parental strains used for GTF single or GTF/Pol II double mutant analyses were constructed by chromosomal integration of GTF mutants into their respective native locus by way of two-step integrations [79]. Strains used in ChIP-exo were TAP-tagged [117] at target genes (*SSL2, SUA7*) using homologous recombination of TAP tag amplicons obtained from the yeast TAP-tag collection [118] (Open Biosystems) and transferred into our lab strain background [119]. All strains with mutations at chromosomal loci were verified by selectable marker, PCR genotyping, and sequencing. *rpo21/rpb1* mutants were introduced to parental strains with or without chromosomal GTF locus mutation by plasmid shuffling [120], selecting for cells containing *rpo21/rpb1* mutant plasmids (Leu^+^) in absence of the *RPB1* WT plasmid (Ura^-^), thus generating single *rpo21/rpb1* mutation strain or double mutant strains combining mutations in GTF and *rpo21/rpb1* alleles. Yeast strains in all experiments were grown on YPD (1% yeast extract, 2% peptone, 2% dextrose) medium unless otherwise noted. Mutant plasmids for yeast promoter analyses were constructed by Quikchange mutagenesis (Stratagene) following adaptation for use of Phusion DNA polymerase (NEB) [121]. All oligonucleotides were obtained from IDT. Yeast strains, plasmids, and oligonucleotide sequences are described in **Additional File 1**.

### Sample preparation for 5′-RNA sequencing

Yeast strains were diluted from a saturated overnight YPD culture and grown to mid-log phase (∼1.5×10^7^/ml) in YPD and harvested. Total RNA was extracted by a hot phenol-chloroform method [122], followed by on-column incubation with DNase I to remove DNA (RNeasy Mini kit, Qiagen), and processing with a RiboZero rRNA removal kit (Epicentre/Illumina) to deplete rRNA. To construct the cDNA library, samples were treated with Terminator 5′ phosphate-dependent exonuclease (Epicentre) to remove RNAs with 5′ monophosphate (5′ P) ends, and remaining RNAs were purified using acid phenol/chloroform pH 4.5 (Ambion) and precipitated. Tobacco acid pyrophosphatase (TAP, Epicentre) was added to convert 5′ PPP or capped RNAs to 5′ P RNAs. RNAs were purified using acid phenol/chloroform and a SOLiD 5′ adaptor was ligated to RNAs with 5′ P (this step excludes 5′ OH RNAs), followed by gel size selection of 5′ adaptor ligated RNAs and reverse transcription (SuperScript III RT, Invitrogen) with 3′ random priming. RNase H (Ambion) was added to remove the RNA strand of DNA-RNA duplexes, cDNA was size selected for 90-500 nt lengths. For SOLiD sequencing, these cDNA libraries were amplified using SOLiD total RNA-seq kit (Applied Biosystems) and SOLiD Barcoding kit (Applied Biosystems), final DNA was gel size selected for 160-300 nt length, and sequenced by SOLiD (Applied Biosystems) as described previously [123, 124].

### 5′-RNA sequencing data analyses

SOLiD TSS raw data for libraries 446-465 was based on 35 nt short reads. The data were delivered in XSQ format and subsequently converted into Color Space csfasta format. Raw data for libraries VV497-520 were in FASTQ format. Multiple read files from each library were concatenated and aligned to *S. cerevisiae* R64-1-1 (SacCer3) reference genome from Saccharomyces Genome Database. We explored the possibility that alignments might be affected by miscalling of 5′ end base of the SOLiD reads. We trimmed one base at the 5′ end of the reads of the TSS libraries VV497-520, and aligned the trimmed reads independently from the raw reads for direct comparison. The alignment rates did not differ significantly, indicating 5′ end of our SOLiD libraries reads were not enriched for sequencing errors more than the rest of the reads. Sequences were with Bowtie [125] allowing 2 mismatches but only retaining uniquely mapped alignments. The aligned BAM files were converted to bedgraphs, and 5′ base (start tag) in each aligned read was extracted using Bedtools (v2.25.0) for downstream analyses [126]. Mapping statistics for TSS-seq, MNase-seq, and ChIP-exo libraries are described in **Additional File 2**.

To assess the correlation between biological replicates and different mutants, base-by-base coverage correlation between libraries was calculated for all bases genome-wide and for bases up and downstream of the promoter windows identified by [14](408 nt total width, described below). Given that Pearson correlation is sensitive to variability at lower coverage levels, correlations for positions above a threshold of ≥3 reads in each library. Heat scatter plots were generated the LSD R package (4.0-0) and compiled in Adobe Photoshop. Heatmaps were generated using Morpheus (https://software.broadinstitute.org/morpheus/) or JavaTreeView [127] and Cluster [128].

To create base-by-base coverage in selected windows of interest, computeMatrix reference-point() function from the deepTools package (2.1.0) was used [129]. There were two types of windows of interest. First, the promoter windows were established by expanding 200 nt up and downstream from the TATA/TATA-like elements identified by [14] (here we term them TATA/TATA-like centered windows) (408 nt total width). Most of these windows (5945/6044) were centered on TATA/TATA-like element annotated in [14], while 99 promoters did not have annotated TATA/TATA-like element and were centered on the TFIIB ChIP-exo peak. Second, we established windows centered on transcription start sites (TSSs) to investigate TSSs at promoters in a core promoter element-independent manner (here we term them TSS-anchored windows). For the TSS-anchored windows, we first determined the 50th percentile (median) TSS (see next paragraph for details) in the TATA/TATA-like centered promoter windows with WT TSS reads derived from *RPB1* WT libraries VV446, VV456, VV497, and VV499 (see below) and expanded 200 nt upstream and 200 nt downstream from this “median” TSS position (401 nt total width), adjusting this window one time based on new TSSs potentially present after shifting the window, and then displaying 250 nt upstream and 150 nt downstream from the median TSS position.

Several characteristics of TSS utilization were calculated as following: (1) The position of the TSS containing the 50th percentile of reads in the window and was termed the “median” TSS. (2) Distance between 10th percentile and 90th percentile TSS position in each promoter was used to measure the width of the TSS distribution, termed the “TSS Spread”. Specifically, TSS positions with 10th and 90th percentile reads were determined in a directional fashion (from upstream to downstream), the absolute value of the difference between two positions by subtraction was calculated as “TSS Spread”. (3) Total reads in windows of interest were summed as a measurement of apparent expression. (4) Normalized densities in windows were calculated as fraction of reads at each TSS position relative to the total number of reads in the window. The normalized densities were subsequently used for examination of TSS usage distribution at each promoter independent of expression level, comparison among different libraries, and start site usage pattern changes in mutants, and visualization. We observed that replicates of each strain (WT or mutant) were highly correlated at the base coverage level as well as primary characteristics of TSS usage (distance to core promoter element, apparent expression) as independently shown by pairwise Pearson correlation and Principal Component Analysis (PCA) (prcomp() in R). We therefore aggregated the counts from replicate strains for downstream analyses (*i.e.*, aligned reads for all replicates of each strain were combined and treated as single “merged library”). Mutant vs WT relative changes of median TSS (**Figure 1E**), TSS spread and normalized TSS densities (**Figure 2**) in the indicated windows were calculated in R and visualized in Morpheus or Graphpad Prism 8. Kruskal-Wallis test was employed to test how many promoters have non-identical distribution in all libraries, as previously described [130], with post hoc Dunn’s test to test how many promoters were significantly shifted in each mutant as compared to WT. Mann Whitney U test was also employed to test how many promoters were significantly shifted in each mutant as compared to WT (*p* < 0.05) for all samples where n≥3 biological replicates.

In the TSS motif analyses, two major characteristics were computed. First was TSS usage defined by the number of reads at each TSS divided by the total number of reads in the promoter window. Second, we calculated TSS efficiency by dividing TSS reads at an individual position by the reads at or downstream of the TSS, as a proxy to estimate how well each TSS gets utilized with regard to the available Pol II (TSS efficiency)[61]. TSS positions with ≥20% efficiency calculated with ≤ 5 reads were excluded (which definitionally are only found at the downstream edges of windows). The corresponding −8, −1, +1 position underlying each TSS (N_−8_N_−1_N_+1_ motif) was extracted by Bedtools getfasta (v2.25.0). Start site motif compilation was done by WebLogo for indicated groups of TSSs. Reads for each N_−8_N_−1_N_+1_ motif of interest were summed, and fraction of the corresponding motif usage in total TSS reads was calculated for each library. Differences of fraction of start site motif usage in WT and mutants were calculated by subtracting the WT usage fraction from that in each mutant.

### Northern blotting and RNA analysis

Northern blotting was performed essentially as described [131]. In brief, 20 µg of yeast total RNA was prepared in Glyoxal sample load dye (Ambion) and separated by 1% agarose gel electrophoresis. RNA was transferred on to membrane by capillary blotting for pre-hybridization. Pre-hybridization solution contained 50% formamide, 10% Dextran sulfate, 5× Denhardt’s solution, 1M NaCl, 50mM Tris-HCl pH7.5, 0.1% SDS, 0.1% sodium pyrophosphate and 500 µg/ml denatured salmon sperm DNA. DNA double-stranded probes were generated by PCR and radiolabeled with ^32^P-dATP using the Decaprime II kit (Ambion) according to the manufacturer’s instructions. Blots were hybridized over night at 42°C and washed twice each in 2× SSC for 15 minutes at 42°C, in 5X SSC with 0.5% SDS for 30 minutes at 65°C, and in 0.2× SSC for 30 minutes at room temperature. Blots were visualized by phosphorimaging (Bio-Rad or GE Healthcare) and quantified using Quantity One (Bio-Rad).

### ChIP-exo sequencing

Yeast cells containing the TAP-epitope [117, 118] were grown to an OD of 0.8 then crosslinked with formaldehyde to a final concentration of 1% for 15 min at room temperature. Crosslinking was quenched with a molar excess of Glycine for 5 minutes at room temperature. Crosslinked cells were pelleted, washed, and then lysed in FA lysis buffer [132] using a chilled (−20°C) beadbeater for 3 minutes. The released nuclei were then pelleted and subsequently resuspended in 600 µl of FA Lysis buffer. The resuspended nuclei were sonicated in a Diagenode Bioruptor Pico for 12 cycles (15 seconds on/30 seconds off). Sonicated chromatin was then incubated overnight on Dynabeads conjugated with rabbit IgG (i5006). ChIP-exo was then performed as previously described [95]. The resulting ChIP-exo libraries were sequenced on a NextSeq 500 in paired end mode. Read 1: 40bp and Read 2: 36bp with dual 8bp indexes. Data were aligned to yeast R64-1-1 with BWA-MEM [133] with low quality reads and PCR duplicates removed by Picard (http://broadinstitute.github.io/picard/) and samtools [134].

### Nucleosome MNase sequencing

Nucleosomal DNAs were prepared by a method described elsewhere [135] with the following modifications. Yeast strains were grown in rich medium (YPD) to mid-log phase (∼1.5×10^7^/ml) and cross-linked with methanol-free formaldehyde (1% final concentration, Polysciences Inc) for 30 min and quenched with 0.25M final concentration of glycine (from 2.5M stock, pH 7). Cells were washed and digested with zymolyase-20T (Sunrise International) (6mg for 500ml culture) for ∼17 min or until ∼90% cells appeared as spheroplasts, followed by MNase (Thermo Fisher Scientific) digestion with different amount of MNase to generate “less” and “more” digested nucleosomes (in general, digests were limited such that at least mono, di, and trinucleosomes were still apparent after agarose gel electrophoresis). Crosslinks on nucleosomes were reversed at 65 °C in the presence of Proteinase K (G-Biosciences) overnight. DNA was extracted by phenol/chloroform, and digested with RNase A (Thermo Fisher Scientific) to remove RNAs. Nucleosomal DNA was separated on 1.5% agarose gels containing SYBR gold dye (Thermo Fisher Scientific) and mono-nucleosome bands were identified and selected under blue light and gel purified (Omega Biotek). Mononucleosomal DNA fragments were sequenced on an Illumina HiSeq 2500 instrument (2×125 paired-end sequencing). Paired-end nucleosome reads were aligned to V64 (SacCer3) reference genome using Bowtie2 [136] allowing 1 mismatch, with only uniquely mapped alignments kept. We used Samtools [134] to extract the alignments to build genome coverage for visualization and start and end position of sequenced DNA fragments. Using the start and end positions of each fragments, fragment length and midpoint position of each fragment were calculated.

Midpoints were analyzed in two main windows of interest. First was median TSS centered window (−250 upstream and +150 downstream based on median TSS position as above). Second, windows were identified based on determined WT +1 nucleosome peak position, as described below using custom scripts (NucSeq v1.0)[137]. Midpoints were assigned to relative coordinates of the window and smoothed using a triweight kernel (75 nt up/downstream total width with a uniform kernel with 5 nt up/downstream width) to get a “smoothed” midpoints profile. The nucleosome peak was called by identifying the local maximum using the smoothed profile. This method enabled us to call a single peak position in ranges of 150 nt windows using the smoothed nucleosome midpoints profiles, thus determining one peak per nucleosome. Average chromosomal coverage (sum of raw midpoints divided by current chromosome length) was calculated for each chromosome as a read threshold per position. The first peak downstream of the median TSS position that had larger than or equal to 20% of chromosomal average coverage and was also within a reasonable position range for a +1 nucleosome was annotated as the +1 nucleosome peak at each promoter (if present). +1 nucleosome peaks were separately identified in two WT libraries (replicates for “less” and “more” digested chromatin), The replicates for “less” digested WT +1 nucleosome peaks showed greater correlation. 500 nt up/downstream of these base positions led to 5660 +1 nucleosome centered 1001 nt wide windows, allowing observation of up to 8 nucleosomes surrounding +1 nucleosomes. Nucleosome midpoints were subsequently assigned to this window using the same method as above. Aggregated nucleosome midpoints analysis was done by sorting the promoters by promoter class, expression level (TSS reads in window) followed by summing the nucleosome midpoint counts at each position in the window. For determination of nucleosome repeat length, we first mapped nucleosome midpoints to windows that span 200nt upstream and 800nt downstream of the determined average +1 nucleosome positions in WT, and subsequently computed autocorrelation by distance to estimate the periodicity of the nucleosome midpoint peak signals. The periodicity of nucleosome signals was first confirmed by the sine wave of autocorrelation function, and the nucleosome repeat length was estimated from the distance of the first non-zero positive peak of autocorrelation function (>0.05). Kernel smoothing (5nt up/downstream width) was applied to the autocorrelation function before peak calling to minimize outlier bias.

## DECLARATIONS

### Ethics approval and consent to participate

Not applicable.

### Consent for publication

Not applicable

### Availability of data and materials

Genomics datasets generated in the current study are available in the NCBI BioProject, under the accession number PRJNA522619. Promoters analyzed in yeast, genomic positions, and attributes (ChIP-exo median positions, +1 nucleosome positions, median TSS positions) are described in **Additional File 3**.

### Authors’ contributions

C.Q. analyzed data, made figures, contributed to writing the manuscript. H.J. initiated project, generated strains, prepared material for TSS-seq, generated material and libraries for MNase-seq, analyzed data, piloted most informatics approaches, and generated outline of the manuscript. I.V. generated libraries for TSS-seq. P.Č. generated strains for ChIP-exo analyses. J.A.L. collaborated with H.J. on nucleosome positioning analyses and generated scripts and code for the analyses. T.Z. provided informatics analysis of TSS-seq data. I.M. constructed strains and performed Northern blotting for promoter variant studies. S.S. initiated informatics analyses for TSS-seq in yeast. P.Č. constructed strains and performed Northern blotting for promoter variant studies. K.H.H. and W.K.M.L. prepared ChIP-exo samples for sequencing. W.K.M.L. processed ChIP-exo sequencing reads. A.M.V. analyzed ChIP-exo data and made figures. R.P.M. and C.D.J consulted on Illumina sequencing strategies and library preparation. S-H.Z. implemented MNase analyses as described in [101]. B.F.P. consulted on ChIP-exo and enabled sequencing of ChIP-exo samples. B.E.N provided funding and consulted on development of TSS-seq for yeast Pol II RNAs. C.D.K conceived the project, guided analyses, made figures, provided funding, and wrote the manuscript. All others read and approved the final manuscript.

## Acknowledgements

Mahmoud Bassal and Kaplan lab members are acknowledged for discussions and comments on the manuscript.

## SUPPLEMENTAL FIGURES AND LEGENDS

**Supplemental Figure 1.**
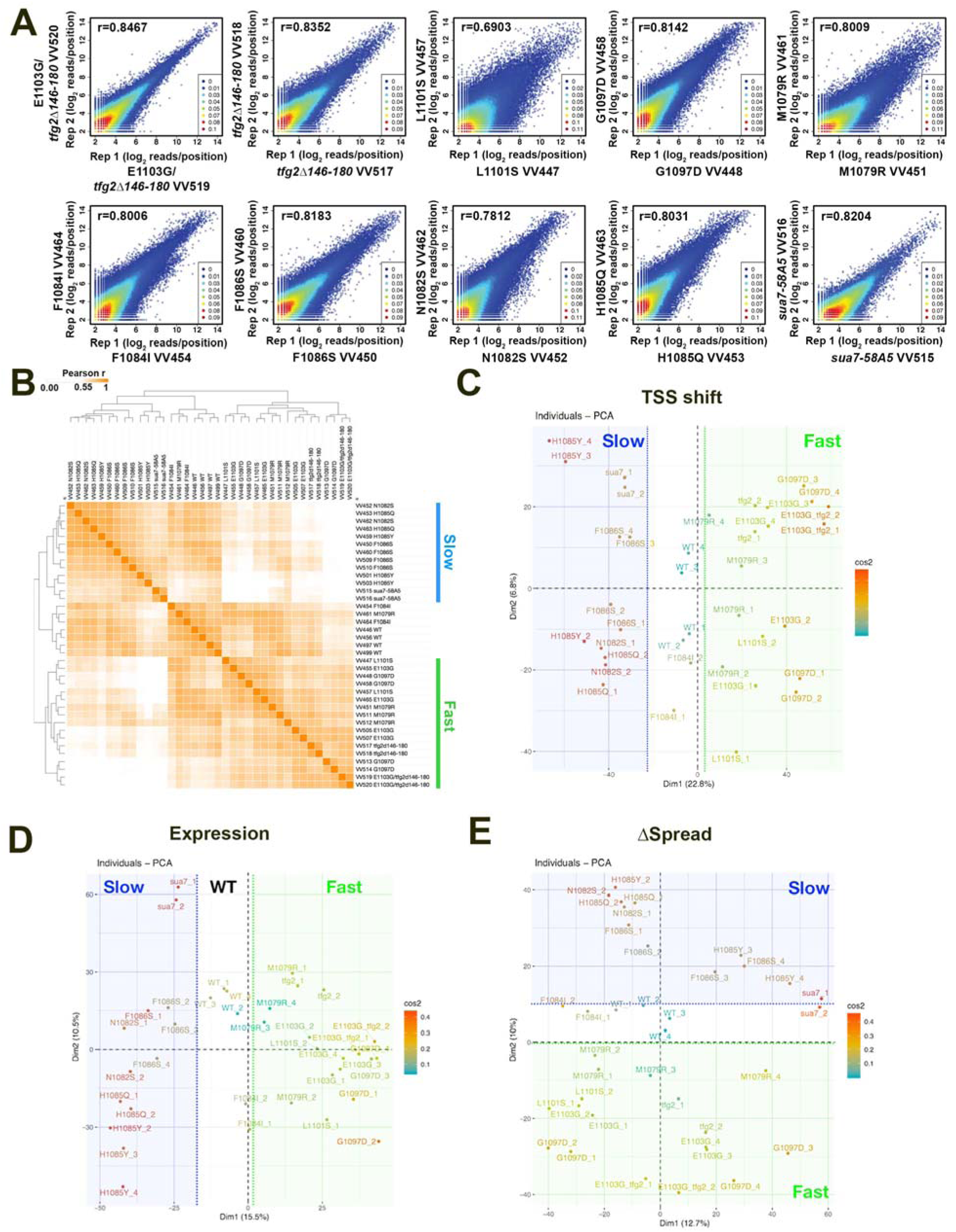
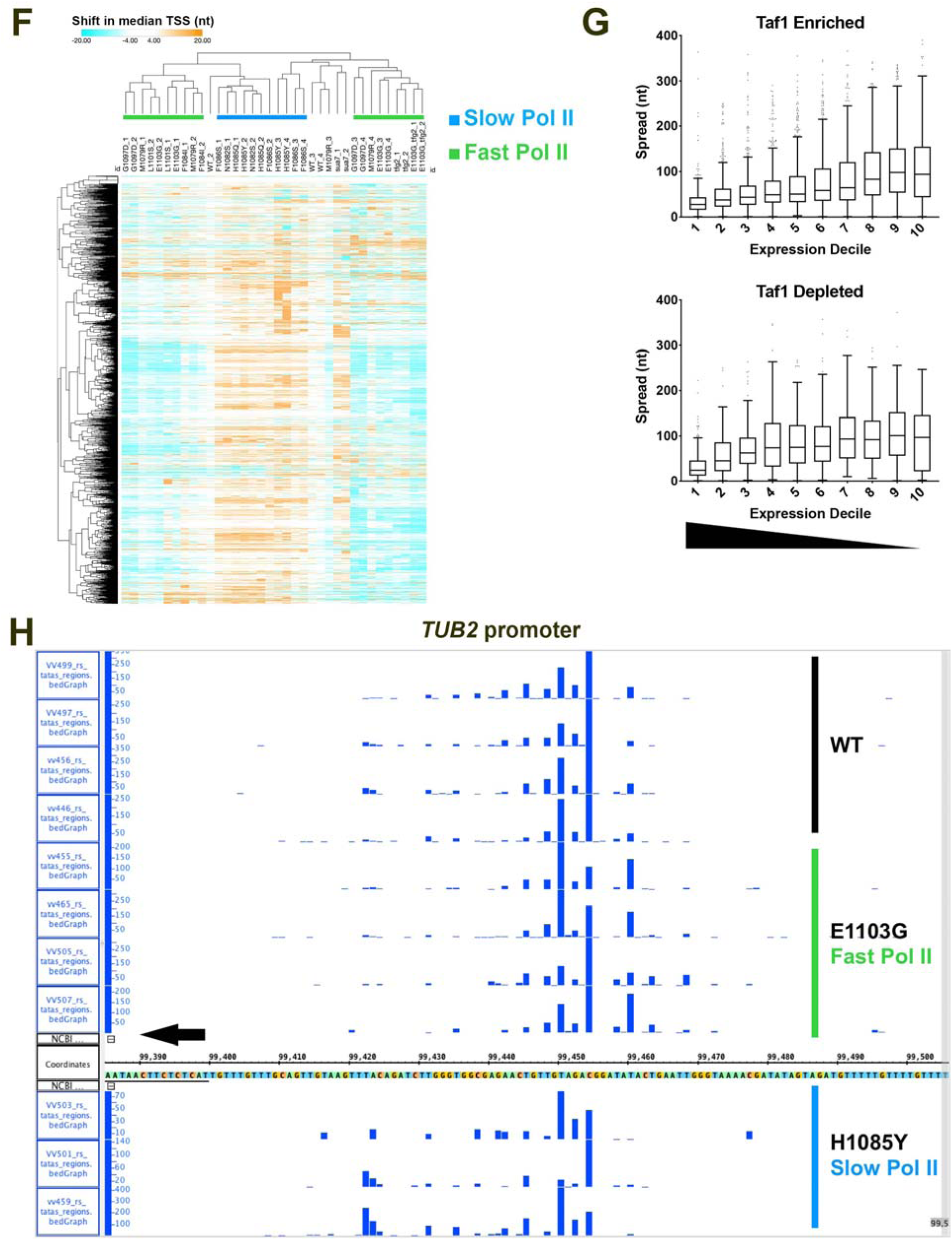
Analysis of TSS-seq replicates. **A.** Example correlation plots for biological replicate TSS-seq libraries. Plots show all genome positions with ≥ 3 reads in each library. Pearson r are listed on each plot. Color scale on the heat scatter plots represent estimated kernel density of the plotted points. **B.** Pearson r correlation coefficients for all TSS-seq library comparisons (examples in A) displayed in a hierarchically clustered heat map. Here, replicates were compared for TSS-seq reads in promoter regions. Promoter regions in this analysis were defined by Rhee and Pugh predicted 8-mer TATA or TATA-like core promoter element position +/- 200 nucleotides upstream and downstream (n=6044). VV numbers represent individual TSS-seq library designations. Libraries VV446-465 represent replicates from one sequencing run (batch one) and VV497-520 represent a separate sequencing run (batch two). Clustering distinguishes two major classes of TSS-seq libraries correlating with upstream TSS-shifting and downstream TSS-shifting mutants. **C.** PCA analysis for individual shifts of TSS medians over all promoters over 200 reads in the aggregated WT data (n=) for individual replicate libraries. Dimension one distinguishes between upstream shifting and downstream shifting initiation mutants across all replicates. Dimension two separates the two sequencing batches (batch two libraries are above *y*=0 and batch one libraries are below *y*=0). **D.** PCA analysis for promoter expression for promoters over 200 reads in the aggregated WT data (n=) for individual replicate libraries. Dimension one distinguishes between upstream shifting and downstream shifting initiation mutants across all replicates. **E.** PCA analysis for Δ TSS Spread for promoters over 200 reads in the aggregated WT data (n=) for individual replicate libraries. Dimension two distinguishes between upstream shifting and downstream shifting initiation mutants across all replicates. Dimension one appears to distinguish primarily between two sequence batches (batch one on left, batch two on right). **F.** Heat map for TSS shifts for individual promoters over 200 reads in the aggregated WT data (n=, *y* axis) determined independently in biological replicate TSS-seq libraries (n=2-4 per strain, *x*-axis). Slow and fast Pol II mutants are distinguished by large bias for downstream (positive, orange) or upstream (negative, cyan) shifts in median TSS position. **G.** Spread of TSSs as determined by the width of the 10-90^th^ percentiles of the TSSs for individual promoters (Taf1 Enriched (n=) or Taf1 Depleted (n=)) inversely correlate with expression. The highest expressed promoters on average have the most focused promoters. Box plots are Tukey plots (see Methods). **H.** Example TSS-seq reads for WT, *rpb1* H1085Y, and *rpb1* E1103G biological replicate TSS-seq libraries at the *TUB2* promoter (n=4,3,4, respectively). The *TUB2* ATG is on the minus strand and designated by the arrow in the lower left.

**Supplemental Figure 2.**
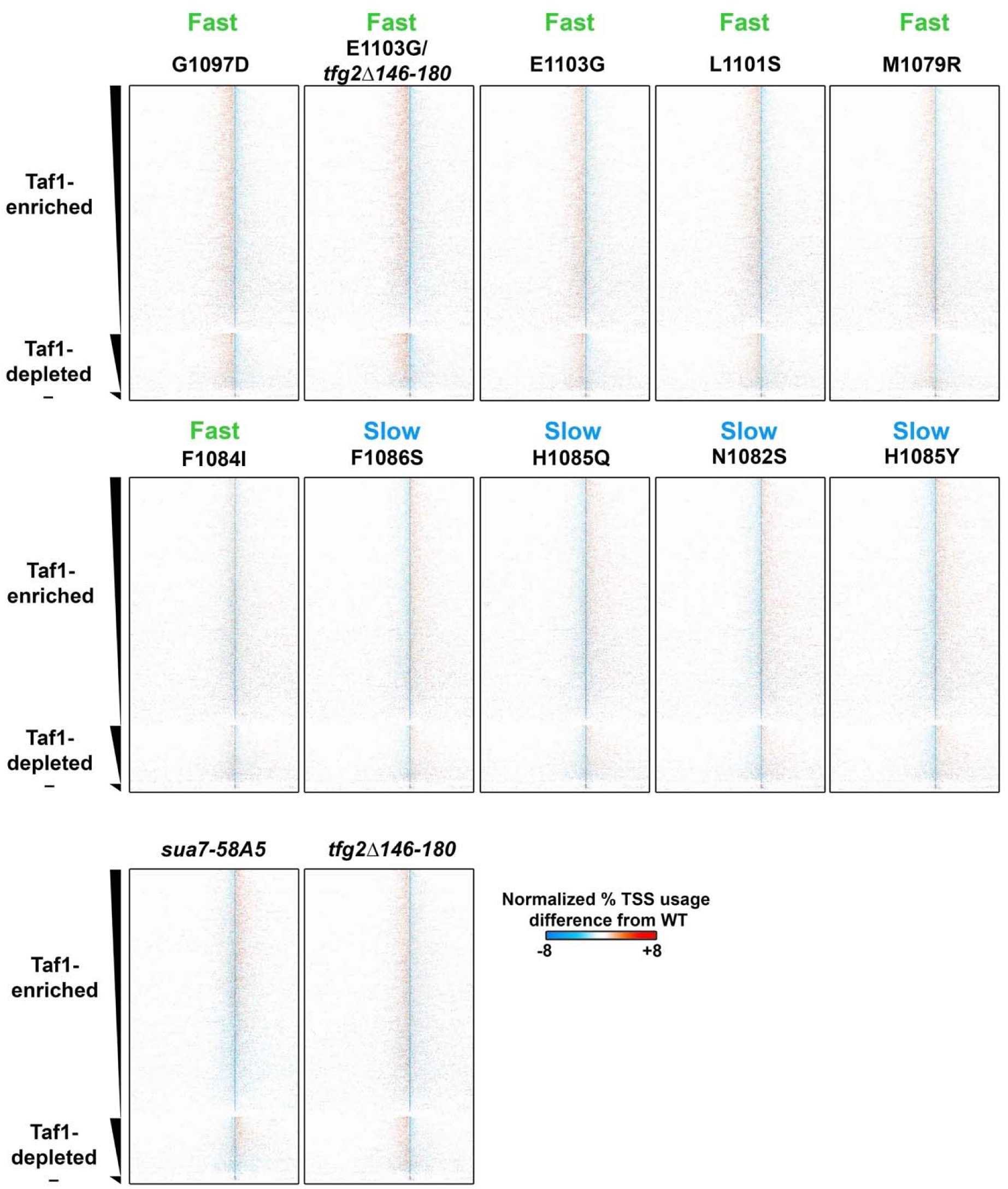
Polar effects on TSS distributions observed for majority of TSS-usage-affecting mutants genome wide. Heat maps as in Figure 2A. *rpb1* H1085Y and *rpb1* E1103G maps from Figure 2A shown here for comparison with all other heat maps. Maps are arranged from strongly upstream shifting to strongly downstream shifting (top left to middle right). Downstream shifting *sua7-58A5* and upstream shifting *tfg2*Δ*146-180* GTF mutants are shown in bottom row.

**Supplemental Figure 3.**
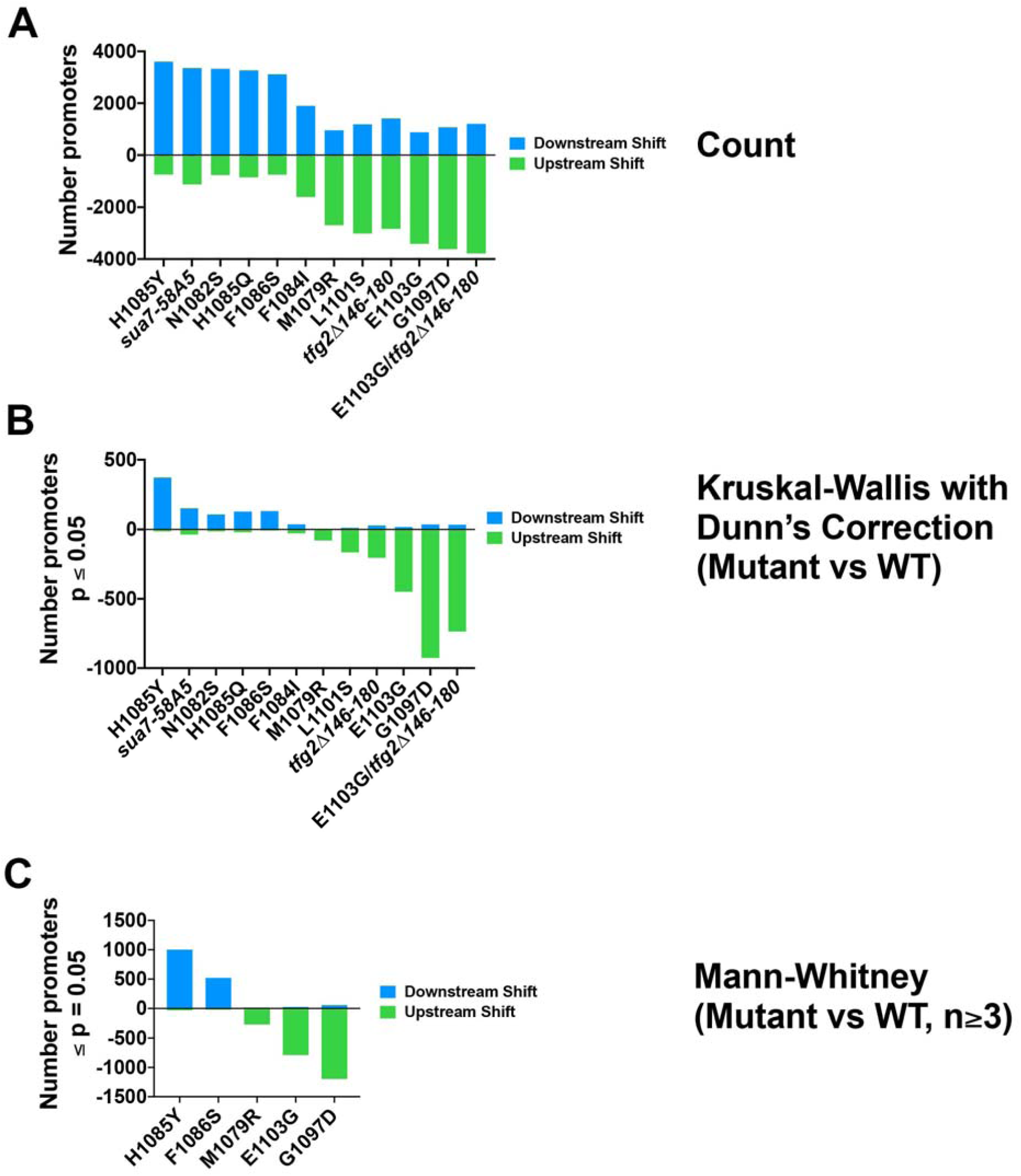
Analysis of promoter level effects of TSS-shifting mutants. **A.** Very large bias in direction of TSS shift depending on mutant class. Counts of promoters with upstream shifts are shown below the *x*-axis (negative numbers) and counts of promoters with downstream shifts are shown above the *x*-axis. **B.** Statistical analysis of significant TSS shifts at individual promoters. TSS shifts for individual promoters across biological replicates for specific mutants were compared to the TSS shifts determined for individual promoters across biological replicates for WT. TSS shifts for each promoter were determined by the median WT TSS position determined from the aggregated WT TSS-seq data. Numbers shown are the individual promoters shifted significantly upstream (negative count) or downstream (positive count) as determined by the Kruskal-Wallis test with Dunn’s correction for multiple comparisons (p≤0.05). The test was performed each promoter, taking into account TSS shift data for all strains (n=2-4). **C**. Mann–Whitney *U* test for TSS shifts at individual promoters as in B, but for each strain where replicates n≥3, which were individually compared with WT (n=4 replicates). Numbers shown are the individual promoters shifted significantly upstream (negative count) or downstream (positive count) in TSS-shifting mutant yeast strains.

**Supplemental Figure 4.**
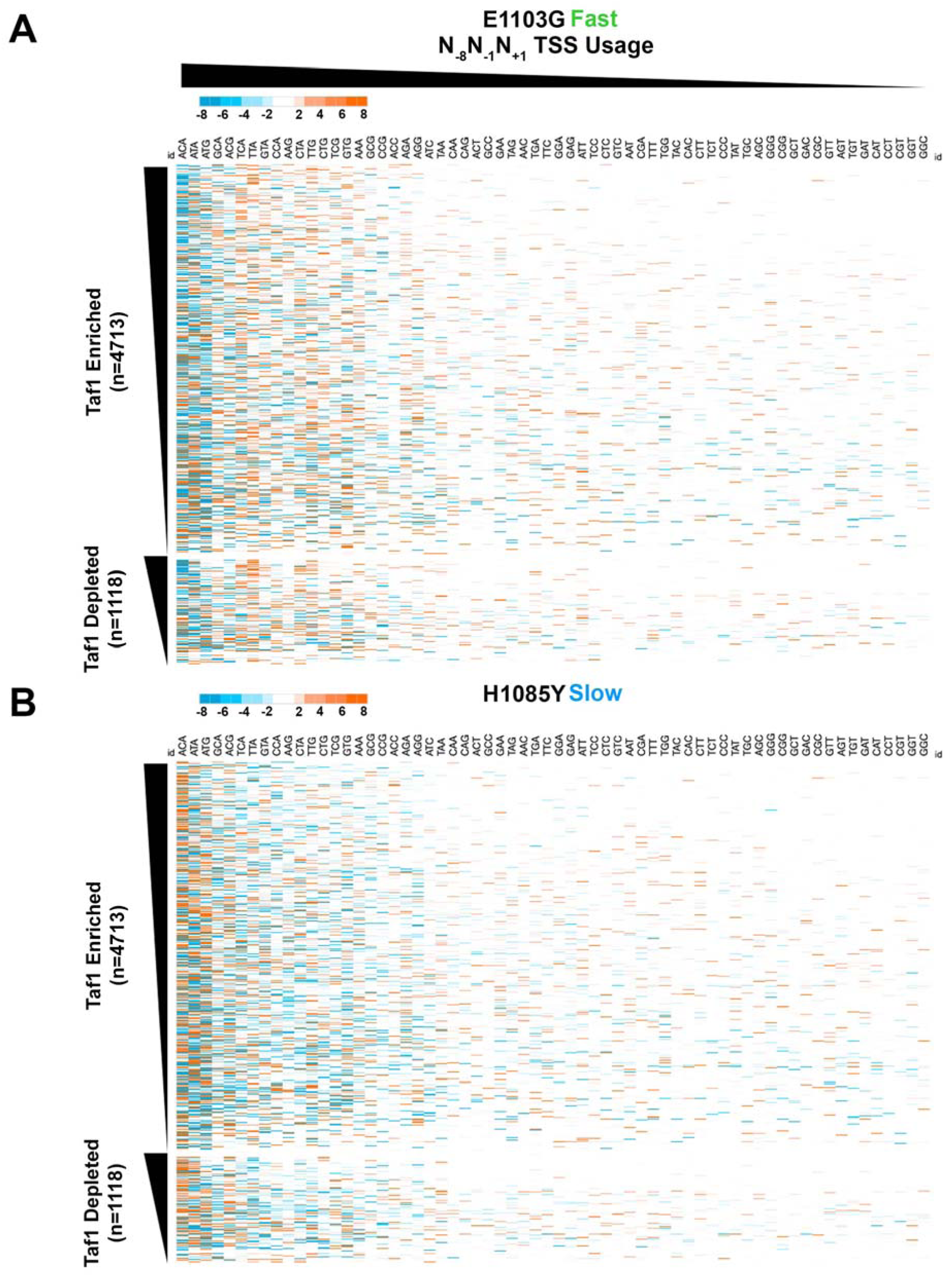
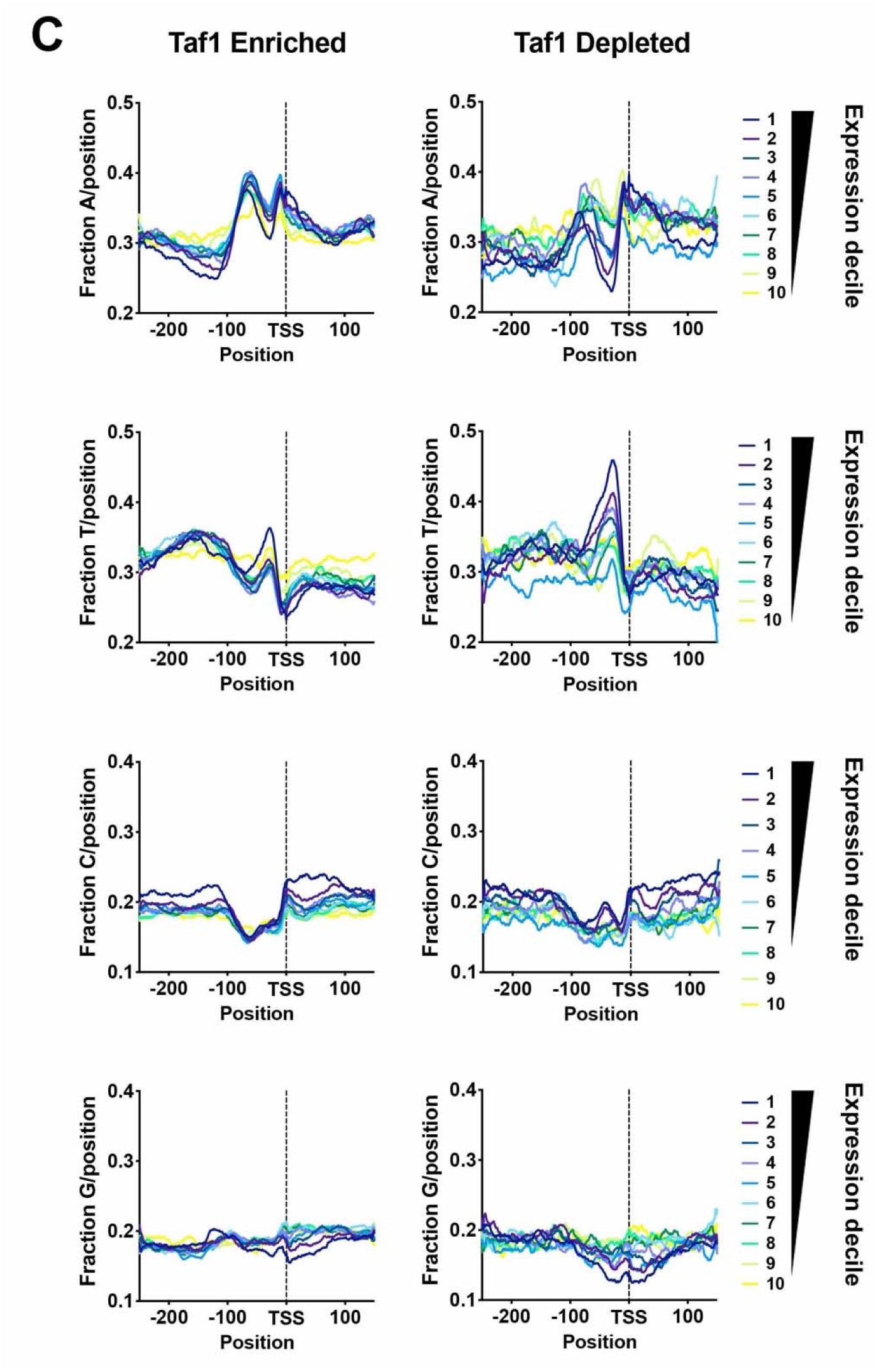
Effects of *rpb1* H1085Y and *rpb1* E1103G mutants on TSS motif usage for N_-8_Y_-1_R_+1_ motifs at the individual promoter level. **A and B.** Heat maps illustrating differences in percent motif usage for individual promoters (*y*-axis) for the 64 N_-8_Y_-1_R_+1_ motifs (*x*-axis) in *rpb1* E1103G (**A**) or *rpb1* H1085Y (**B**) are shown. Motifs are rank ordered based on overall usage across genome in WT yeast (high to low from left to right) and promoters are separated into Taf1 Enriched and Taf1 Depleted classes and rank ordered within class by expression (high to low from top to bottom). **C.** Distribution of bases on the top promoter strand for Taf1 Enriched or Taf1 Depleted promoters, separated by expression decile in WT cells.

**Supplemental Figure 5.**
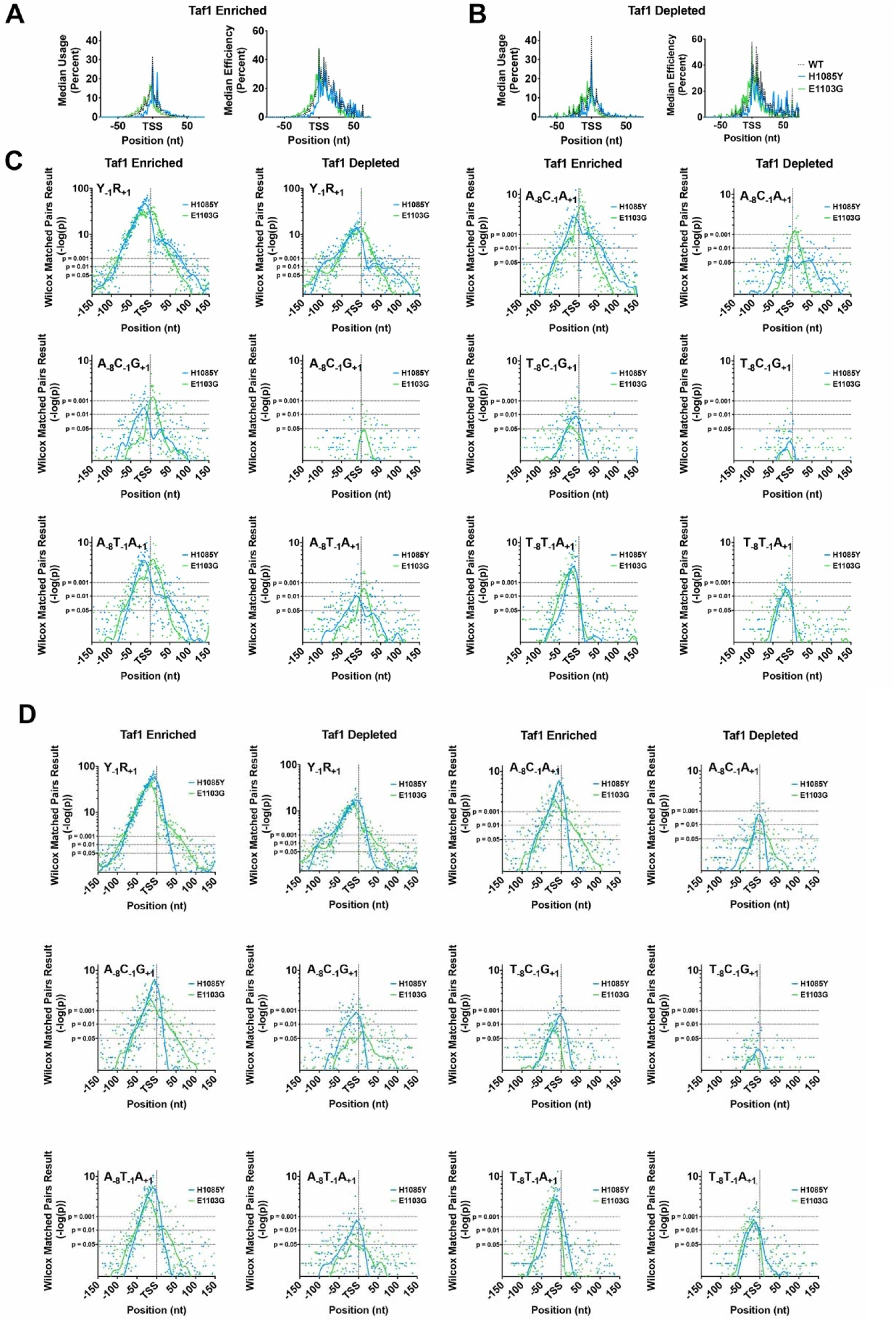
TSS-shifting mutants alter TSS usage and efficiency across TSS motifs throughout promoter regions. **A.** Example median TSS usage (left) for A_-8_C_-1_A_+1_ motifs for WT, *rpb1* E1103G, and *rpb1* H1085Y strains, or TSS efficiency for A_-8_C_-1_A_+1_ motifs for the same strains (right) across Taf1 Enriched promoters. Median usage/efficiency determined from the subset or promoters that have an A_-8_C_-1_A_+1_ motif at the designated promoter position (see schematic in Figure 4C). **B.** Same as (A) but for Taf1 Depleted promoters. **C.** Statistical analysis of distributions of TSS Usage between WT and *rpb1* E1103G, or WT and *rpb1* H1085Y strains for specified motif at each promoter position. The Wilcoxon Matched-Pairs Signed Rank test was used to compare distributions of WT usage of a particular motif for a particular promoter position with the distributions of usage for those motifs/positions in Pol II mutants. Therefore, there is a p-value determined for each promoter position for each motif. The -log(p) of each p-value is displayed on the *y*-axis of a plot for a number of example motifs, separated into Taf1 Enriched and Depleted promoters. The lines are LOWESS smooths of individual points. Note that some positions will have no motifs across promoters at a particular position and significance will in part be determined by how many instances of motif there are at each position across promoters. Almost all motifs shown exhibit clusters of positions with significant differences between mutants and WT. **D.** Analysis as in (C) but for TSS efficiencies.

**Supplemental Figure 6.**
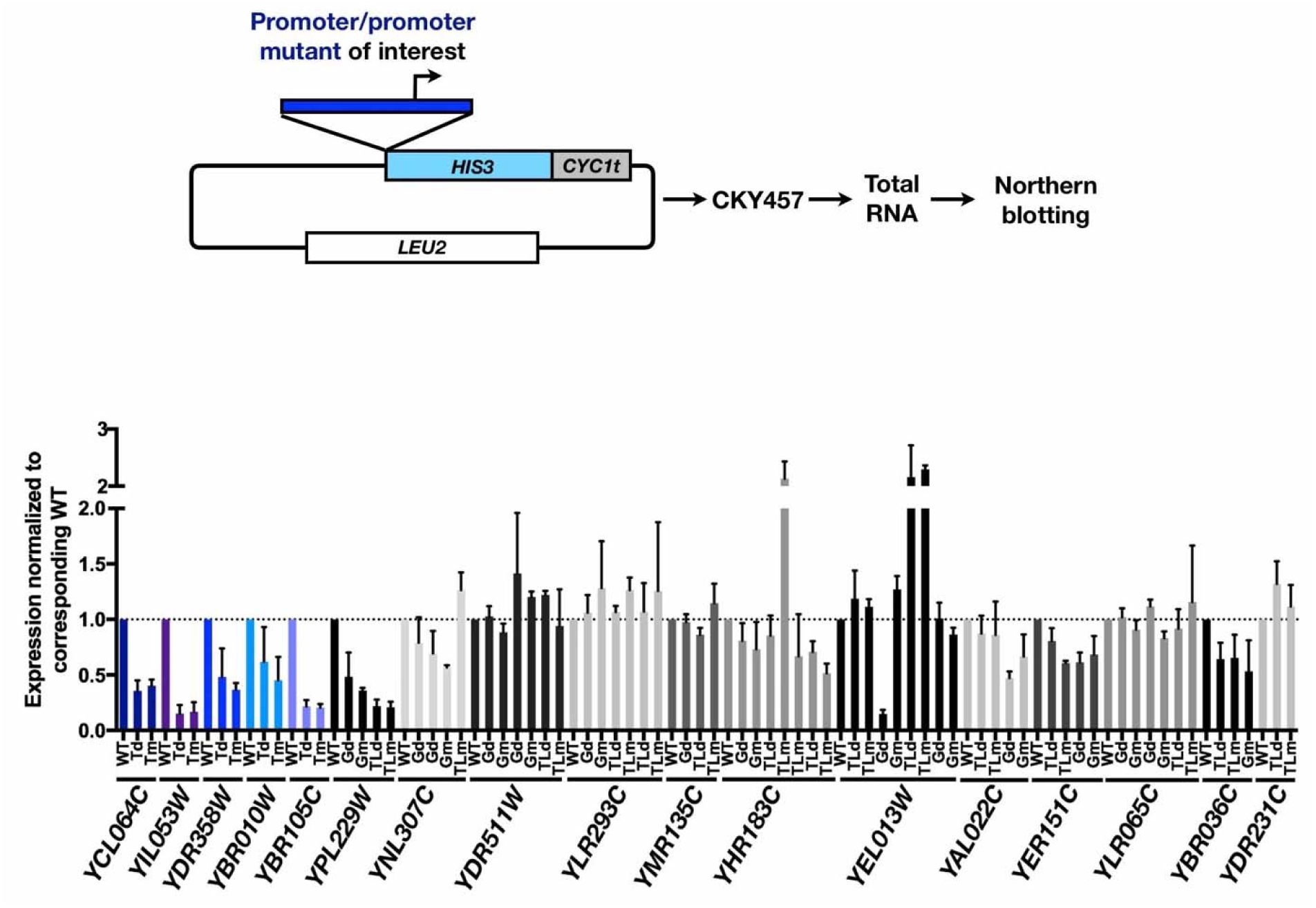
Effects on expression level of putative core promoter element mutations. (Top) Schematic of reporter plasmids fusing promoters of interest (up to ATG) to a *HIS3* ORF/*CYC1* terminator reporter. (Bottom) Quantification of Northern blotting for control WT or promoters mutated (“Tm”) or deleted (“Td”) for consensus TATA elements (promoters shaded in blue), mutated or deleted for GAE (“Gm” or “Gd”, respectively) or mutated or deleted for TATA-like elements identified by Rhee and Pugh or our own analyses (“TLm”, “TLd”, respectively). Bars are mean +/- standard deviation of the mean (n=≥3).

**Supplemental Figure 7.**
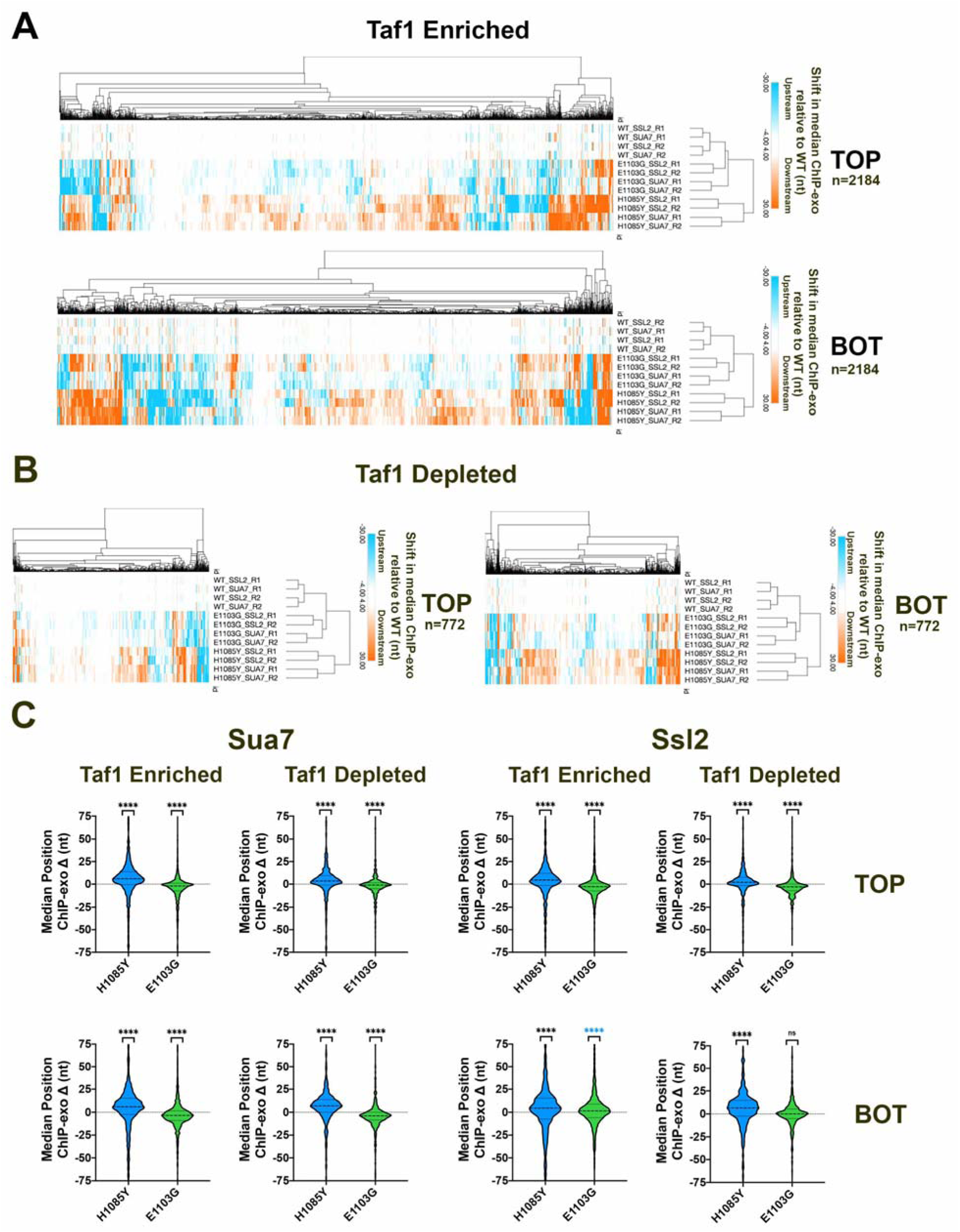
ChIP-exo analysis if Sua7 and Ssl2 in *rpb1* catalytic mutants. **A.** Heat map of median ChIP-exo value shifts relative to WT for individual ChIP-exo replicates for top (TOP) or bottom (BOT) DNA strand signal for Taf1 Enriched promoters representing the top ∼50% of overall ChIP-exo signal as determined by WT signal for each factor. ChIP-exo signal medians were determined for each promoter window on both strands for two biological replicates for each strain. Median positions were determined for individual replicates and the WT average position was subtracted. A positive value (orange) indicates that a mutant has a downstream shift in ChIP-exo signal while a negative value (cyan) indicates an upstream shift. **B.** Same as in (A) but for Taf1 Depleted promoters. **C.** Statistical analysis of average shift in median ChIP-exo signal on top (TOP) or bottom (BOT) DNA strands for Sua7 or Ssl2 at Taf1 Enriched promoters (left) or Taf1 Depleted promoters in WT or *rpb1* mutants. Promoter-mapped ChIP-exo tags compared for two biological replicates for WT, *rpb1* E1103G, and *rpb1* H1085Y in Ssl2-TAP and Sua7-TAP strains. Median of averaged shifts in *rpb1* H1085Y or *rpb1* E1103G compared to zero (no shift) by Wilcoxon Signed Rank Test (**** indicates p<0.0001). Note that Ssl2 for Taf1 Enriched BOT in E1103G shifts downstream 1 nt (marked with blue asterisks) and this is the opposite shift than for all other significant effects.

**Supplemental Figure 8.**
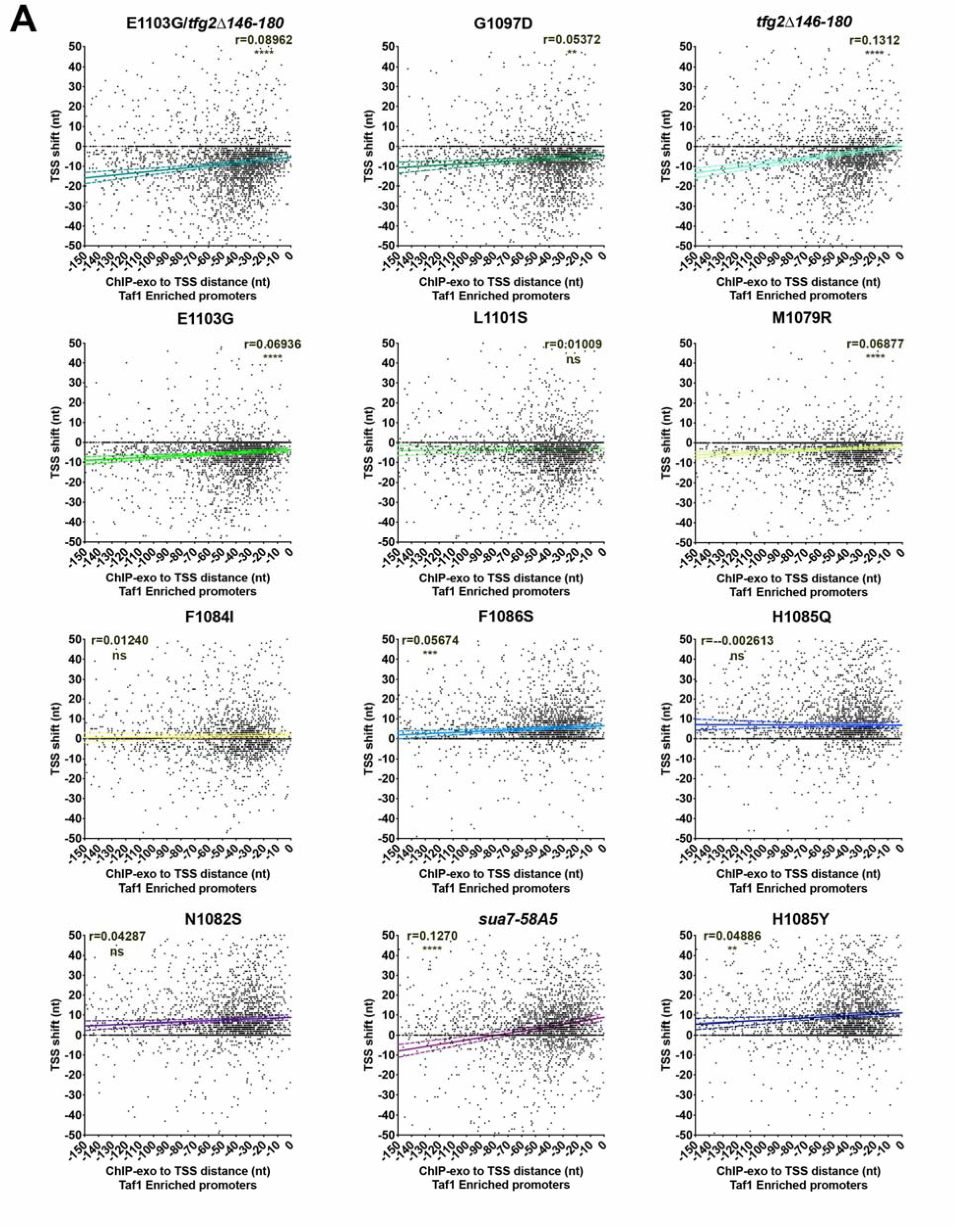
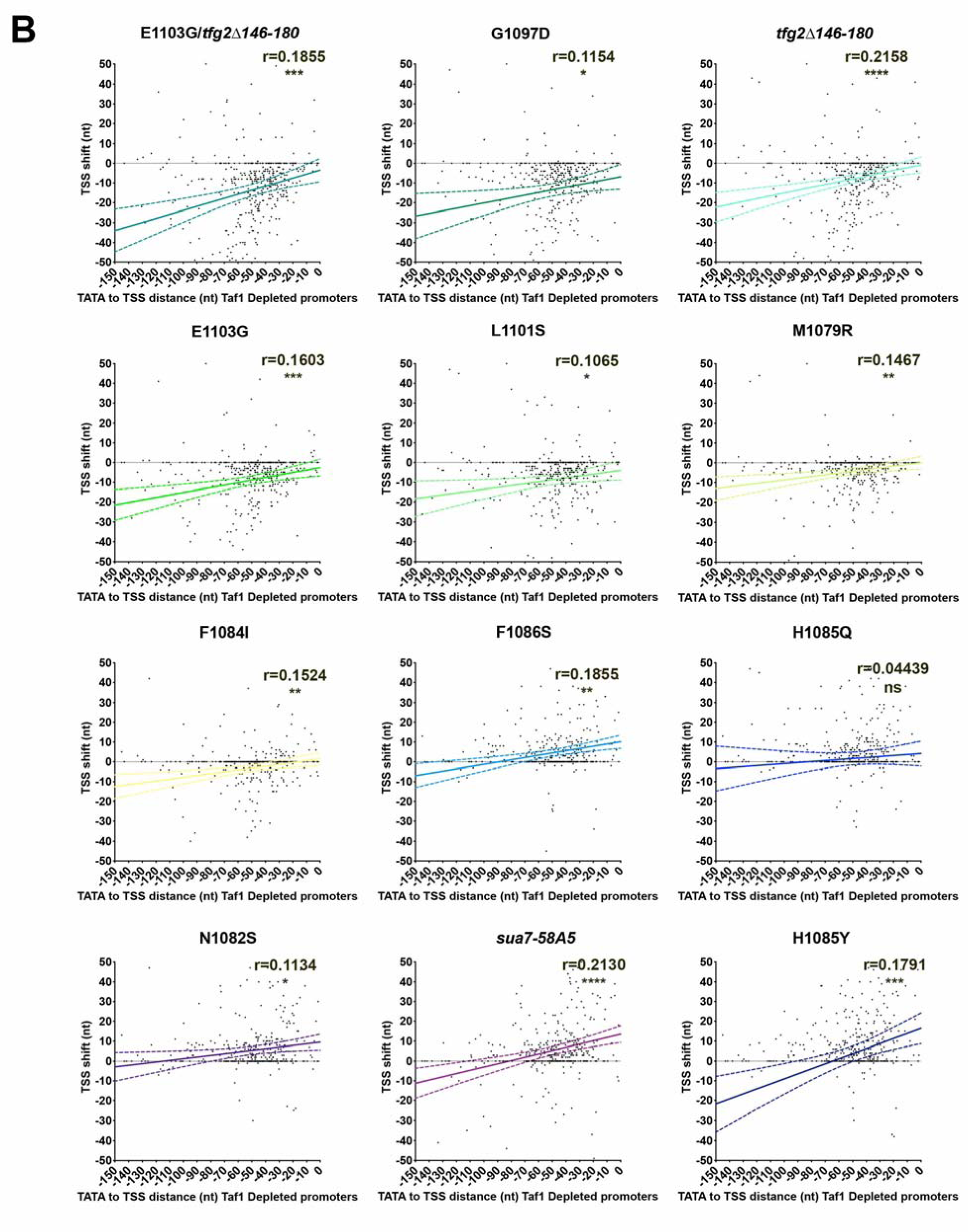
Correlation of TSS shift with ChIP-exo or core promoter element-TSS distance. **A.** Median TSS shifts (*y-*axis) for promoters ≥ 100 reads expression in WT for denoted TSS mutants for Taf1 Enriched promoters plotted versus ChIP-exo-TSS distance (*x*-axis). Lines are linear regression with 95% confidence interval for the linear fit. Pearson r correlations are shown for each plot with asterisks indicating P value (two-tailed, (0.0332 (*), 0.0021 (**), 0.0002 (***), <0.0001 (****)). **B.** Median TSS shifts (*y-*axis) for promoters ≥ 100 reads expression in WT for denoted TSS mutants for Taf1 Depleted promoters with consensus TATA elements plotted versus consensus TATA-TSS distance (*x*-axis). Lines are linear regression with 95% confidence interval for the linear fit. Pearson r correlations are shown for each plot with asterisks indicating P value (two-tailed, (0.0332 (*), 0.0021 (**), 0.0002 (***), <0.0001 (****)).

**Supplemental Figure 9.**
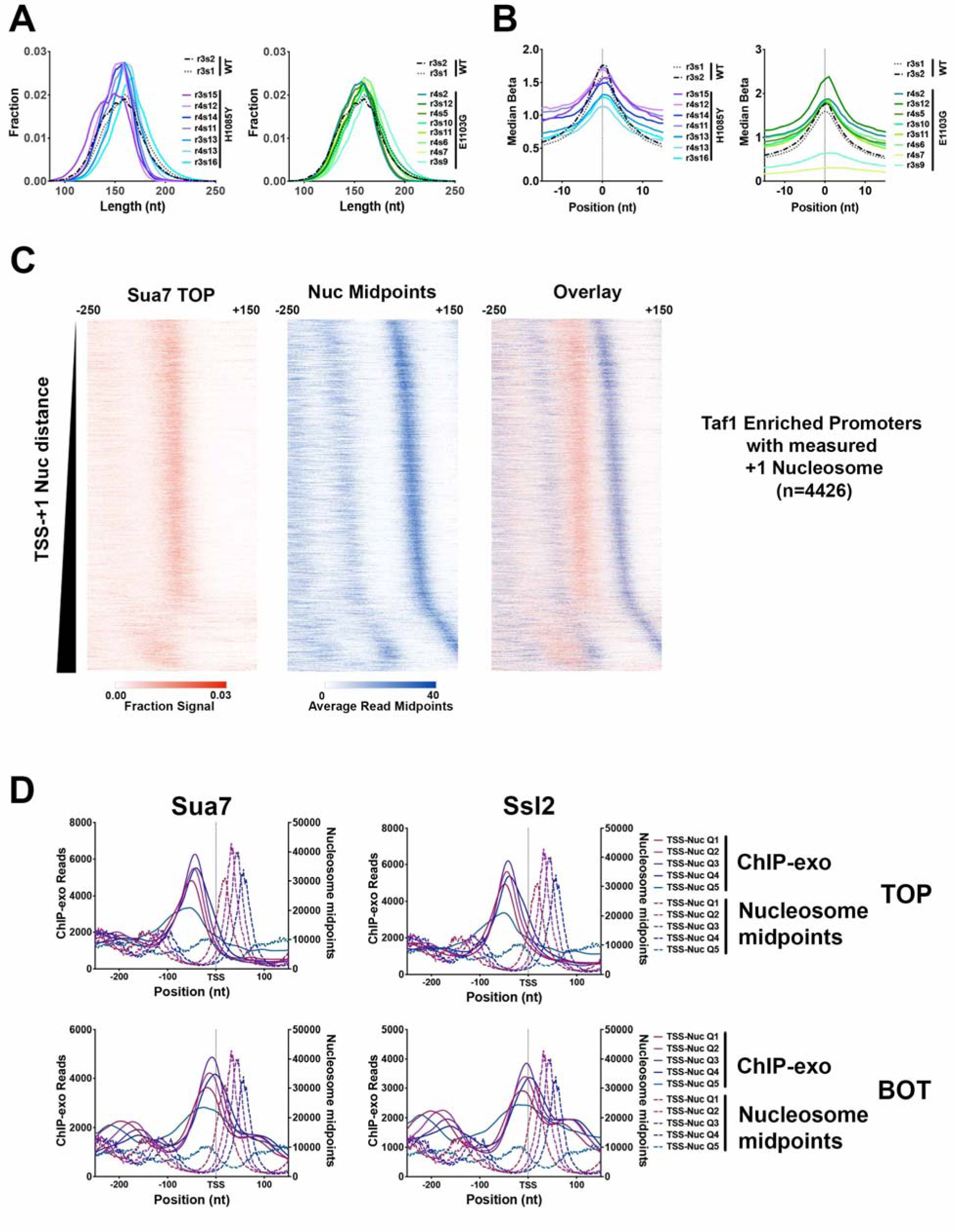
MNase-seq analyses of nucleosome positions in WT, *rpb1* H1085Y, and *rpb1* E1103G mutants. **A.** Paired-end sequencing fragment length distributions in WT and H1085Y MNase-seq libraries (left) and in WT (as left, shown for reference) and E1103G MNase-seq libraries (right). Libraries arranged within groups from most digested (top) to least digested (bottom). **B.** Probability of nucleosome positioning (“Beta”) values determined by method of Zhou *et al* for MNase-seq libraries arranged as in (A). **C.** Heat maps of Sua7 TOP strand ChIP-exo signal (right), nucleosome +1 midpoints (middle) or overlay of the two (left) indicating correlation of PIC component localization and nucleosome positioning for Taf1 Enriched promoters. **D.** Nucleosome midpoints as determined by MNase-seq (dashed lines) and GTF ChiP-exo signals for Taf1 Enriched promoters (solid line LOWESS smooth of scatter plots) were aggregated by promoter quintiles determined by TSS-+1 nucleosome midpoint position. Nucleosome midpoints are from WT strain and the same data are shown as reference for each ChiP-exo plot. First to fifth quintiles are promoters with the closest +1 nucleosome to furthest, respectively. Fifth quintile promoters likely have a weak +1 nucleosome and thus the determined +1 nucleosome is in some cases like the +2. ChiP-exo aggregate data shows intermediate correlation with +1 nucleosome-TSS distance.

**Supplemental Figure 10.**
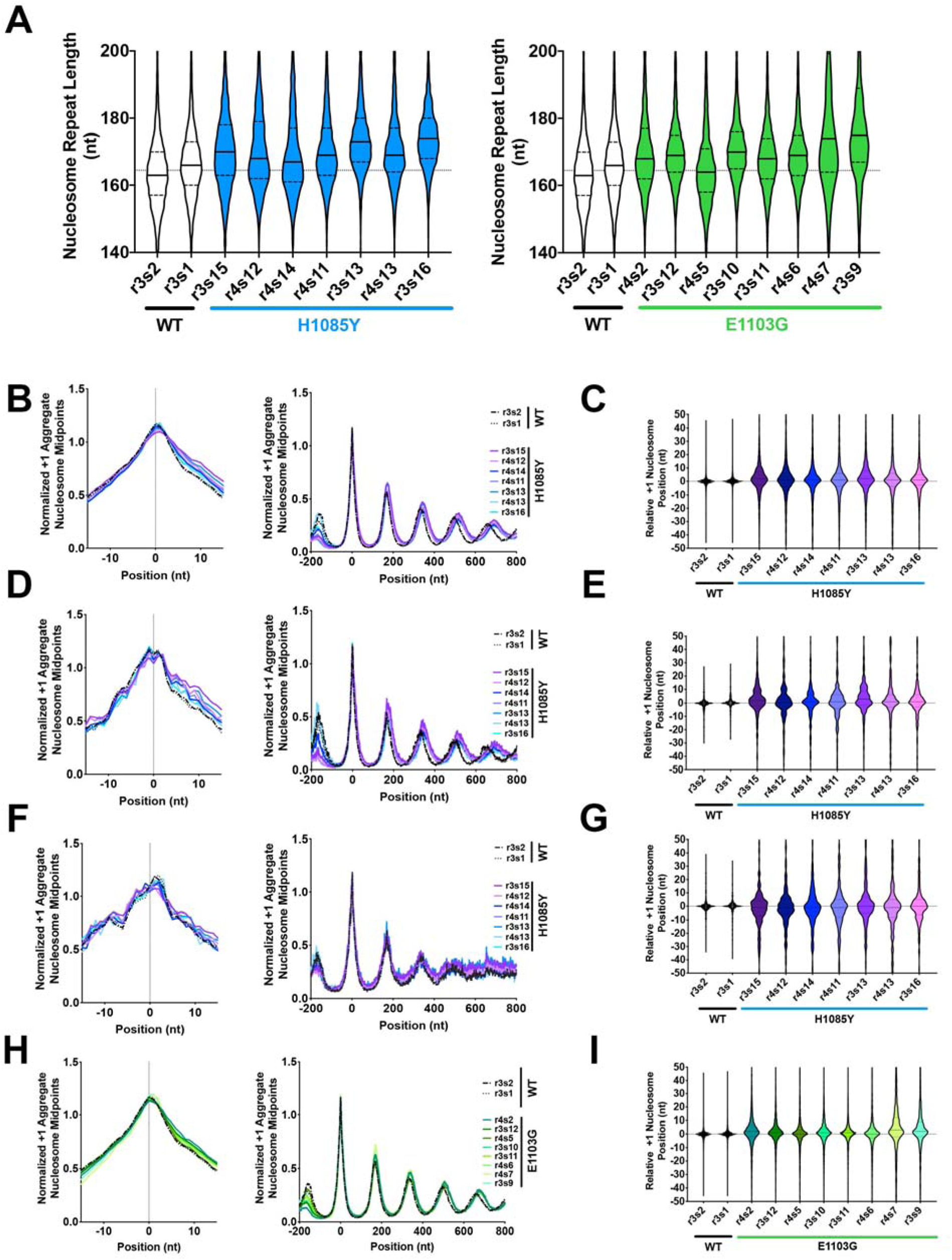
Relationship of promoter chromatin architecture to PIC position and effects of TSS-usage affecting mutants on nucleosome positioning. **A. (Left)** WT MNase-seq replicates (n=2) compared to *rpb1* H1085Y MNase-seq replicates (n=7) for nucleosome repeat length as determined by autocorrelation analysis (see Methods). **(Right)** Same as left but for *rpb1* E1103G vs WT (WT samples same on left). **B.** Nucleosome positioning in WT and *rpb1* H1085Y for Taf1 Enriched promoters aligned by +1 nucleosome in WT (left), over genes (−200 to +800 from +1 nucleosome position, right). **C.** Determined +1 nucleosome position for WT and *rpb1* H1085Y Taf1 Enriched promoters for individual MNase-seq libraries relative to position determined by averaging the four WT libraries. Box plots are Tukey plots (see Methods). **D. and E.** Nucleosome positioning analyses as in (B, C) for top expression decile Taf1 Enriched promoters for WT and *rpb1* H1085Y. **F. and G.** Nucleosome positioning analyses as in (B, C) for bottom expression decile Taf1-enriched promoters for WT and *rpb1* H1085Y. **H. and I.** Nucleosome positioning analyses as in (B, C) for Taf1 Enriched promoters for *rpb1* E1103G. WT data from (B, C) shown as reference.

